# Simultaneous single-cell CRISPR, RNA, and ATAC-seq enables multiomic CRISPR screens to identify gene regulatory relationships

**DOI:** 10.1101/2025.02.11.637716

**Authors:** Kaivalya Shevade, Yeqing Angela Yang, Kevin Feng, Karl Mader, Volkan Sevim, Jacob Parsons, Gunisha Arora, Hasnaa Elfawy, Rachel Mace, Scot Federman, Rustam Esanov, Shawn Shafer, Eric D Chow, Laralynne Przybyla

## Abstract

The ability to identify gene functions and interactions in specific cellular contexts has been greatly enabled by functional genomics technologies. CRISPR-based genetic screens have proven invaluable in elucidating gene function in mammalian cells. Single-cell functional genomics methods, such as Perturb-seq and Spear-ATAC, have made it possible to achieve high-throughput mapping of the functional effects of gene perturbations by profiling transcriptomes and DNA accessibility, respectively. Combining single-cell chromatin accessibility and transcriptomic data via multiomic approaches has facilitated the discovery of novel cis and gene regulatory interactions. However, pseudobulk readouts from cell populations can often cloud the interpretation of results due to a heterogeneous response from cells receiving the same genetic perturbation, which could be mitigated by using transcriptional profiles of single cells to subset the ATAC-seq data. Existing methods to capture CRISPR guide RNAs to simultaneously assess the impact of genetic perturbations on RNA and ATAC profiles require either cloning of gRNA libraries in specialized vectors or implementing complex protocols with multiple rounds of barcoding. Here, we introduce CAT-ATAC, a technique that adds CRISPR gRNA capture to the existing 10X Genomics Multiome assay, generating paired transcriptome, chromatin accessibility and perturbation identity data from the same individual cells. We demonstrate up to 77% guide capture efficiency for both arrayed and pooled delivery of lentiviral gRNAs in induced pluripotent stem cells (iPSCs) and cancer cell lines. This capability allows us to construct gene regulatory networks (GRNs) in cells under drug and genetic perturbations. By applying CAT-ATAC, we were able to identify a GRN associated with dasatinib resistance, indirectly activated by the HIC2 gene. Using loss of function experiments, we further validated that the gene, ZFPM2, a component of the predicted GRN, also contributes to dasatinib resistance. CAT-ATAC can thus be used to generate high-content multidimensional genotype-phenotype maps to reveal novel gene and cellular interactions and functions.

## Introduction

Since the discovery of CRISPR-Cas9 genome editing over a decade ago, researchers have been adopting the system in a wide range of applications due to its high efficiency and ability to target specific genomic loci. CRISPR-mediated perturbations (knockout, interference, activation) provide a powerful tool to interrogate gene functions in a systematic manner. Given its ease of programmability, CRISPR-Cas9 can be combined with genome-wide guide RNA (gRNA) libraries for conducting unbiased and high-throughput genetic screens^1^. Different phenotypic assays can be utilized to evaluate the outcome of a pooled CRISPR screen, including gRNA representation analysis and single-cell evaluation^2,3,4^. By using single-cell RNA-seq (scRNA-seq) as a readout for pooled CRISPR screens, methods such as Perturb-seq^5,6^, CRISP-seq^7^, and CROP-seq^8^ allow us to dive deep into the relationship between every genetic perturbation and its corresponding transcriptomic outcomes. These functional genomic studies provide information-rich phenotypic profiling that can significantly accelerate our understanding of biological and pathological processes.

In addition to transcriptomic changes, epigenomic effects are often as informative and crucial to understanding transcriptional regulation in a complex system. The dynamics of chromatin accessibility, dictated by transcription factor occupancy and epigenetic mechanisms such as histone modifications and DNA methylation, underlie many fundamental biological processes in development and disease^9,10^. Thus, single-cell ATAC-seq (Assay for Transposase-Accessible Chromatin with sequencing)^11^, which measures chromatin accessibility in individual cells, has been implemented as a readout for pooled CRISPR screens, and is represented by methods such as Perturb-ATAC^12^, CRISPR-sciATAC^13^, and Spear-ATAC^14^. These methods employ various single-cell platforms and have guide capture efficiencies ranging from 40% to 90%. As an example of their utility, recently published atlases detailing single-cell chromatin accessibility in various tissues offer a abundant insights into cell type-specific cis-regulatory elements and how they can be used to predict links between non-coding disease risk variants to downstream affected genes and functions^15,16^.

Single-cell profiling techniques such as these can provide novel insights into gene regulatory networks that drive several complex biological processes, with applications to developmental trajectories and disease mechanisms. Drug resistance to chemotherapeutics and/or targeted therapies is a known problem in cancer and can account for almost 90% of cancer-associated deaths^17^. Chronic myeloid leukemia (CML) develops from a recombination between chromosomes 9 and 22 creating a short chromosome 22 (Philadelphia chromosome) in myeloid cells that results in a BCR-ABL gene fusion, causing constitutive activation of the Abl tyrosine kinase. Dasatinib is an FDA-approved drug for the treatment of CML. Dasatinib, like most other treatments for CML, is a broad-spectrum tyrosine kinase inhibitor (TKI)^18^. The development of TKIs has increased 5-year survival rate from 22% in the mid 1970s to 69%^19^. However, there is no complete cure for the disease. Although dasatinib and other TKIs effectively block the Abl kinase, drug resistance ultimately leads to treatment failure. In as many as 40% of cases of clinical TKI failure, cancer cells exhibit sustained BCR-ABL inhibition^20^ but instead activate alternative pathways which help them survive in presence of the drug. Identification of BCR-ABL-independent mechanisms of drug resistance could lead to new therapeutic targets for CML^21,22^. Therefore, understanding the dynamics of transcriptome and DNA accessibility in response to genetic and drug perturbation might help us piece together the novel gene regulatory relationships that underlie drug resistance by providing a deeper understanding of the direct and indirect effects of alternative pathway activation.

To interrogate both the transcriptional profile as well as the epigenetic landscape in CRISPR screens, we have developed CAT-ATAC (<u>C</u>RISPR <u>A</u>nd <u>T</u>ranscriptomics-<u>A</u>ssay for <u>T</u>ransposase-<u>A</u>ccessible <u>C</u>hromatin). This method simultaneously profiles the transcriptome, chromatin accessibility, and CRISPR gRNA expression within single cells by leveraging the Multiome assay kit from 10X Genomics and is compatible with the widely used Perturb-seq dual-guide vector backbone as well as other gRNA vectors. The pairing of the two layers of data, transcriptional and regulatory states, at single-cell resolution provides a multidimensional phenotypic readout that will enable CRISPR-based screens in a range of applications intended to uncover gene-regulatory networks. In this study, we developed CAT-ATAC in iPSCs and used it to elucidate a novel gene regulatory network that underlies dasatinib drug resistance in CML cells.

## Results

### CAT-ATAC methodology and proof-of-concept

CAT-ATAC is built on the 10X Genomics Multiome assay, which uses a droplet-based method to capture both RNA and chromatin fragments. In the workflow (Figure 1A), dual-guide direct capture constructs^23^ are introduced into cells that express Cas9-CRISPR machinery (eg, CRISPRn, CRISPRi, or CRISPRa). CAT-ATAC reagents are combined with tagmented nuclei and passed through the 10X Genomics single-cell controller to compartmentalize each nucleus into an oil droplet containing a mix of reagents, enzymes and oligo primers. Through downstream thermocycling reactions and enrichment steps, the mRNA transcripts, ATAC fragments, and the gRNA transcripts within each cell are barcoded with a unique sequence and constructed into three libraries, one each for assaying RNA, ATAC and gRNA. Figure 1B details the steps involved in capturing gRNAs, which involves the spike-in of a pre-annealed reverse transcription (RT) primer with a splint oligo to reverse transcribe the gRNA. The splint oligo is complementary to the spacer sequence on the ATAC-capture oligo of the multiome gel beads and serves as a bridge between the downstream gRNA RT products and the gel bead. The RT primer contains a sequence complementary to the capture sequence (CS1) found in gRNAs expressed from the dual-guide Perturb-seq gRNA construct^24^, a UMI, and the Nextera R1 sequence and forms a duplex with the splint oligo through a partial Nextera R1 sequence. During RT and template switching, the spiked-in primer initiates first strand cDNA synthesis of the gRNA followed by template switching. Subsequent rounds of PCR amplification add appropriate adaptors and sample indexes to make the final single-cell gRNA library.

**Figure 1:**
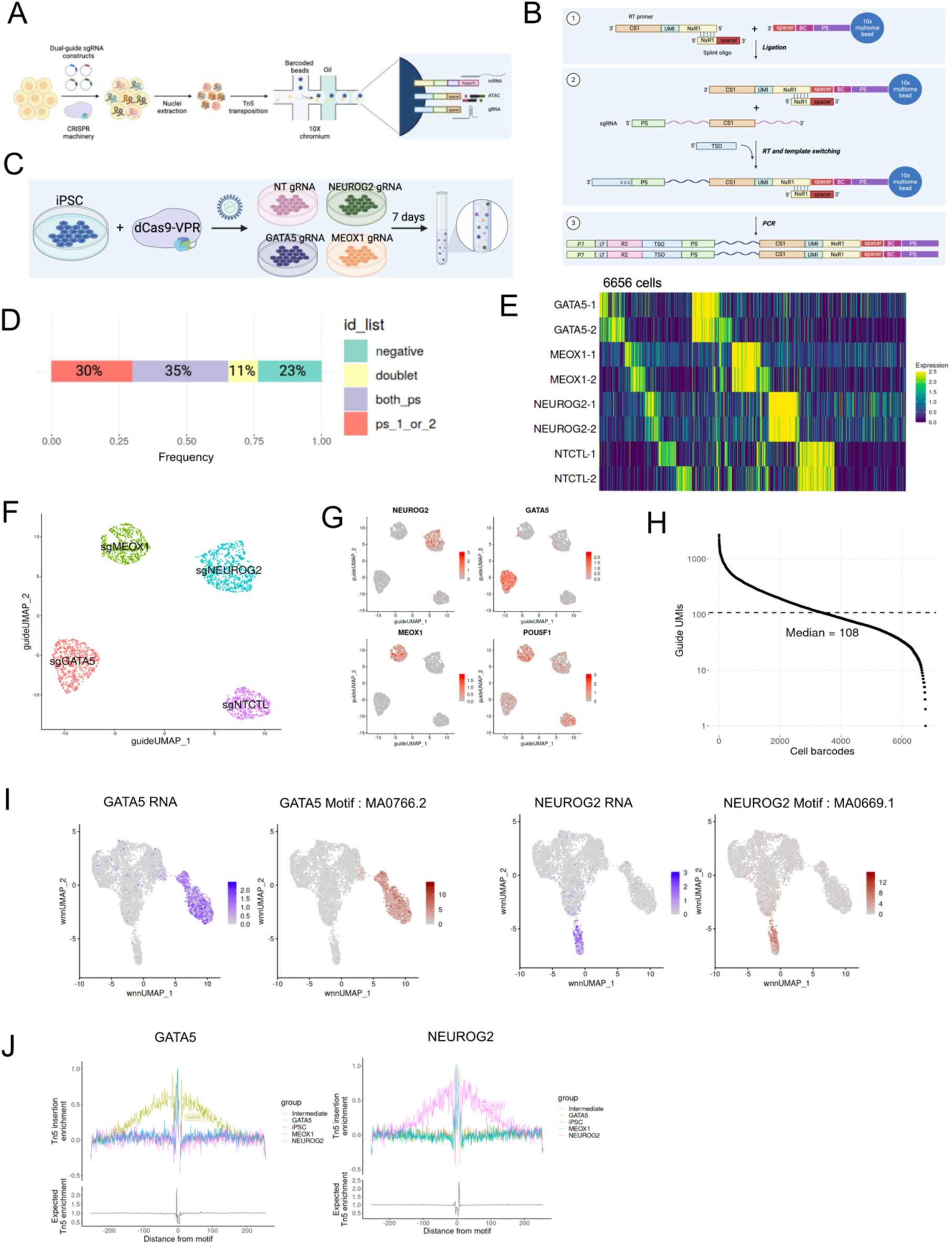
CAT-ATACmethod and validation A. Illustration of CAT-ATAC workflow B. Detailed schematic illustrating the guide capture process. C. Pilot experiment overview D. Overall guide assignment for the pilot experiment. 76% of the total cells could be assigned to guides, 11% of which were identified as doublets. E. Heatmap showing guide assignment in all 6656 cells. F. UMAP embedding of singlets expressing 2 protospacers (Singlet_2_PS) based on guide expression. G. RNA expression plots of targeted genes in guide expression UMAP. H. Barcode rank plot for sgRNA UMI counts, showing a median of 108 UMIs for captured guide sequences per cell. I. Expression of the TF shown in purple, motif enrichment shown in brown. J. TF footprinting for GATA5 and NEUROG2.

For proof-of-concept, we performed a pilot experiment (Figure 1C). We used a human induced pluripotent stem cell (iPSC) line engineered with dox-inducible CRISPRa-VPR and delivered 4 dual gRNA constructs into the cells by lentiviral transduction in an arrayed format. Each dual gRNA construct expresses 2 different activating guide sequences for the same target, in this case, 3 transcription factors (NEUROG2, GATA5, MEOX1) and a non-targeting control (NT); guide sequences were selected from the top two performing guides in the Weissman V2 CRISPRa library as listed in Supplemental Table 1. We activated transcription factors in the pilot experiment to induce directed differentiation through overexpression of lineage-specific transcription factors^25^, based in part on prior work that established sets of key regulators leading to directed differentiation in iPSCs^26,27^. NEUROG2 is a well-established driver of neuronal cell fate^28^, whereas GATA5 and MEOX1 overexpression caused loss of pluripotency markers in iPSCs^26^ but led to an undefined cell fate. By turning on the expression of these TFs in iPSCs through CRISPR-mediated activation, we expect the cells to exhibit changes in RNA and ATAC profiles that are distinct from those observed in the parental iPSCs, making this an ideal system to evaluate our CAT-ATAC method. After dox induction of CRISPR activation targeting these transcription factors in iPSCs for 7 days, the cells were pooled and analyzed by CAT-ATAC. It is worth noting that excess cells were cryopreserved and used in subsequent iterations for optimization and no loss in performance was observed, demonstrating the flexibility and utility of this assay.

We obtained a total of 6,656 cells with both RNA and ATAC profiles after data preprocessing and filtering, of which 23% of cells could not be assigned to a gRNA identity, 30% of cells were assigned to a single gRNA, 35% of cells were assigned to a pair of protospacers (PS) with the same target, and 11% were classified as doublets which contained more than one gRNA with incongruous targets that could be from droplets containing multiple cells (Figure 1D). The heatmap in Figure 1E depicts guide expression in all 6,656 cells with their gRNA identity labeled in each row. Upon filtering for only cells expressing paired gRNAs (n = 2,361 cells), we plotted the guide classification in a UMAP embedding (Figure 1F) and observed 4 distinct clusters corresponding to the 4 groups of cells receiving a different guide construct, indicating faithful gRNA identification. Figure 1G shows the RNA expression of the target genes on the guide UMAP with successful overexpression due to CRISPR-mediated gene activation. POU5F1 (gene name for OCT4) is a canonical iPSC marker, and its expression can be seen in the negative control gRNA cluster as well as the MEOX1 cluster but is reduced in NEUROG2 and GATA5. Violin plots in Supplemental Figure 1A further demonstrate that TF upregulation is only observed in the cluster receiving the corresponding gRNAs, and POU5F1 is downregulated in both the GATA5 and NEUROG2 group, but not the MEOX1 group, suggesting that unlike cells in the former two groups, MEOX1 cells were not fully differentiated and may not yet have exited their pluripotent state, or may be on a lineage trajectory that requires sustained expression of POU5F1. Guide count rank plot in Figure 1H shows a median of 108 guide UMIs per cell. This is lower than the reported guide UMI of ∼1000 in Perturb-seq^24^ due to differences in the assay chemistries and cell lines used but is sufficient to assign guide identity to a majority of the cells with high confidence, supported by the target gene expression data for each gRNA cluster.

Next, we proceeded with further evaluations of the pilot CAT-ATAC data by integrating both the RNA and ATAC profiles for each cell. Shown in Supplemental Figure 1B, we performed clustering and constructed UMAP embeddings with just the RNA, ATAC, or weighted nearest neighbor (WNN) analysis^29^. We classified the cells into 5 clusters, including one for each of the overexpressed TF, one for iPSCs, and another termed as intermediate due to the lack of expression of iPSC markers or targeted TFs. Motif enrichment was carried out for each cluster using Signac ^30,31^, and we observed a high level of correlation between RNA expression of the TF (in purple) and enrichment of its motif (in brown) (Figure 1I). Similarly, transcription factor footprinting analysis indicates increased Tn5 accessibility flanking the motif locations (Figure 1J). We compared CAT-ATAC to other recently published single-cell RNA-seq, ATAC-seq and joint multiome technologies and found better performance of RNA and ATAC capture, including increased RNA UMIs and genes per cell (Supplemental Figure 1C and D), as well as increased ATAC fragments and peaks per cell (Supplemental Figure 1E and F). This indicated that CAT-ATAC can successfully be used to analyze perturbation-specific effects on RNA and ATAC profiles of cells.

### Genome-wide CRISPRi screen identifies gene knockdowns that affect sensitivity to dasatinib

Next, we sought to integrate and apply the RNA and ATAC data to identify how genes affected by specific perturbations drive the mechanisms of cancer drug resistance and identify targets that can disrupt the transcriptional programs responsible. For this application, we targeted CML resistance mechanisms to TKIs that are driven by activation of pathways independent of BCR-ABL mutations. We performed a genome-wide CRISPRi screen to identify genetic modifiers of sensitivity to the TKI dasatinib and hypothesized that some of these genetic modifiers must control transcriptional programs driving resistance.

The screen was performed in K562 cells, a CML cell line that harbors a BCR-ABL mutation, and the experimental design is outlined in Supplemental Figure 2A. After observing high concordance between the replicates (Figure 2A) we identified 164 dasatinib sensitizing gene knockdowns and 800 dasatinib de-sensitizing gene knockdowns that were differentially enriched using an adjusted p-value cutoff of 0.05 and a phenotype score cutoff of +/-0.1 (Figure 2B). These include previously reported known TKI sensitizing genes BCL2L1, MCL1 and PI3K-AKT pathway genes as well as desensitizing genes BAX and LZTR1 (Figure 2B and Supplemental Figure 2B)^32,21,33^. We decided to focus on sensitizing gene hits as we hypothesized that some of these genes might constitute the cellular response to dasatinib and potentially be responsible for resistance due to aberrant pathway activation. We filtered down the list of 164 sensitizing gene hits to 60 hits by first removing 61 genes which had significant growth phenotype scores in our vehicle control sample or which were identified as a core essential gene by Hart et. al. ^34^ and by keeping only the genes which have nuclear function to focus on potential core components of gene regulatory networks (Supplemental Figure 2C). We performed arrayed validation for these 60 gene hits by transducing K562 cells expressing CRISPRi machinery with an arrayed dual-guide library for the 60 genes and 2 non-targeting controls such that each vector targeting one gene had the 2 top performing guides from the pooled survival screen (Figure 2C). Following FACS analysis we calculated percent increase in cell death and confirmed that all the gene knockdowns sensitized cells to dasatinib when compared to NTCs (Figure 2D). GO analysis indicated that these 60 genes broadly included functional classification of transcriptional regulation, protein ubiquitination and RNA binding (Figure 2E). Of the 20 genes with transcriptional control functions there were 7 genes with known transcriptional repression function, 3 genes with known transcriptional activator function and 10 genes with unknown transcriptional activity. The transcriptional networks controlled by these 20 genes could regulate cellular resistance to dasatinib and reveal new potential therapeutic targets. We hypothesized the global changes to the transcriptomic and DNA accessibility profiles after knockdown would shed light on the underlying regulatory networks and effectors that confer resistance to dasatinib. Hence, we used CAT-ATAC to evaluate the effect of gene perturbations simultaneously on transcriptome and DNA accessibility from the same cells.

**Figure 2:**
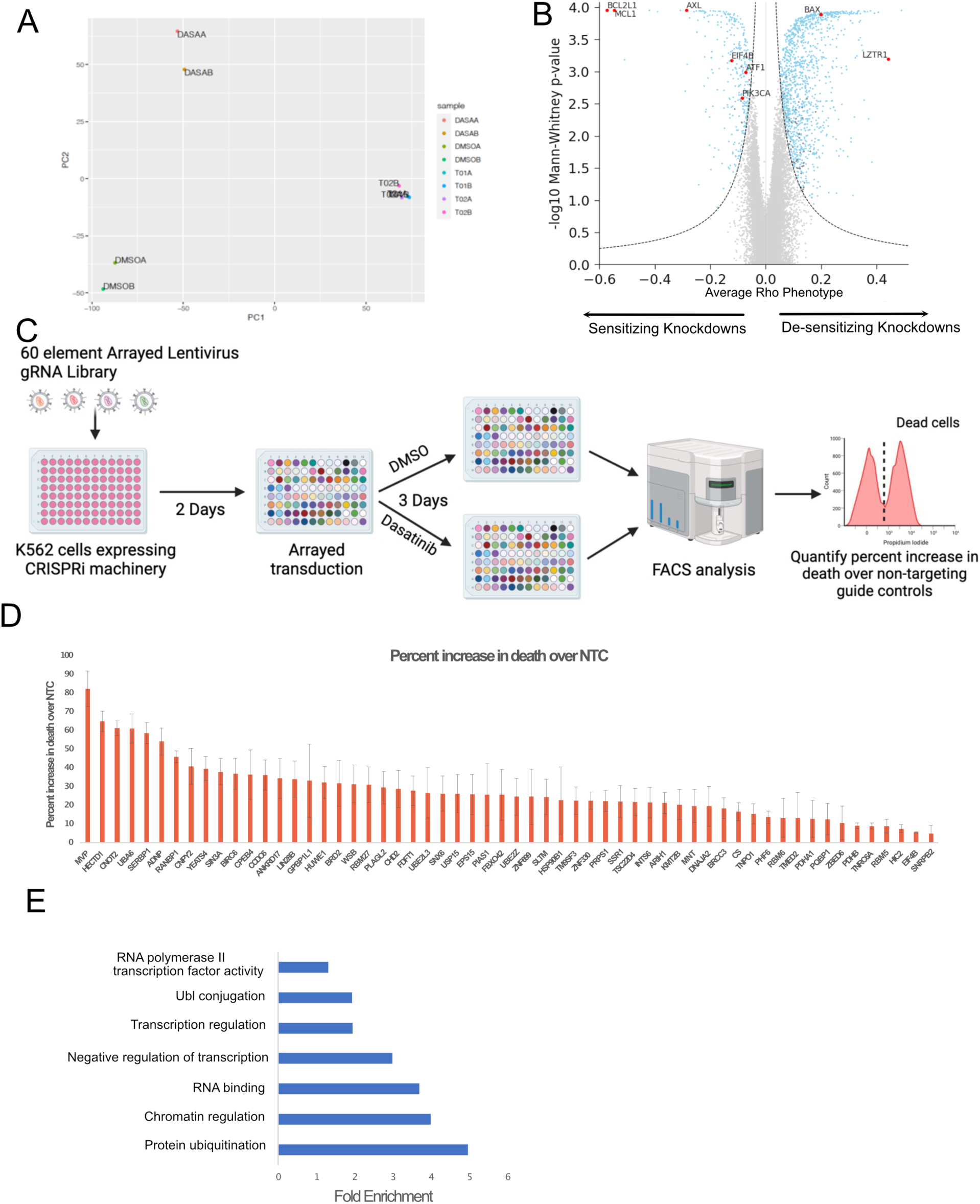
Pooled CRISPRi screen identifies gene knockdowns which modulate resistance to dasatinib A. PCA plot showing reproducibility of replicate samples from the genome-wide screen. B. Volcano plot showing gene knockdowns exhibiting sensitizing and desensitizing phenotypes. Blue dots represent all significant hits. Genes highlighted in red were previously known to affect dasatinib sensitivity. C. Schematic showing overview of the arrayed validation experiment conducted to validate hits from the primary genome-wide screen. D. Bar plot displaying results from the arrayed validation experiment showing percent increase in cell death for each of the guide expressing cells over non-targeting control guide expressing cells. Error bars represent standard deviation. E. GO analysis for the 60 genes included in the arrayed validation experiment.

### Identification of perturbation-specific effects on dasatinib-induced alterations to DNA accessibility and transcription

K562 cells expressing CRISPRi machinery were transduced with a pooled lentiviral library containing 26 pairs of gRNAs. The 26-element library comprised 20 dual-guide vectors targeting the 20 transcriptional regulatory genes identified above and 6 non-targeting gRNA pairs. The gRNA sequences used are listed in Supplemental Table 1. We harvested the cells 48hr after dasatinib treatment to minimize loss due to cell death and investigate the earliest transcriptional and DNA accessibility changes that happen upstream of cell death (Figure 3A). The experimental design consisted of 2 biological replicates resulting in 4 pooled CAT-ATAC samples, two treated with dasatinib and two treated with DMSO as a vehicle control. We processed the 4 samples individually using the Seurat^30^ analysis package and achieved 72-77% efficiency of guide assignment across the 4 samples (Supplemental Figure 3A). The UMAP shows high concordance between the replicates (Figure 3B). After guide assignment and filtering, we observed significant knockdowns (median KD 85.12% +/-10% SD) for all the targeted genes except for those where the percentage of cells expressing the target gene was very low (Figure 3C). Concomitantly we saw a decrease in promoter accessibility for many of the targeted genes based on the scATAC data (Figure 3D and Supplemental Figure 3B).

**Figure 3:**
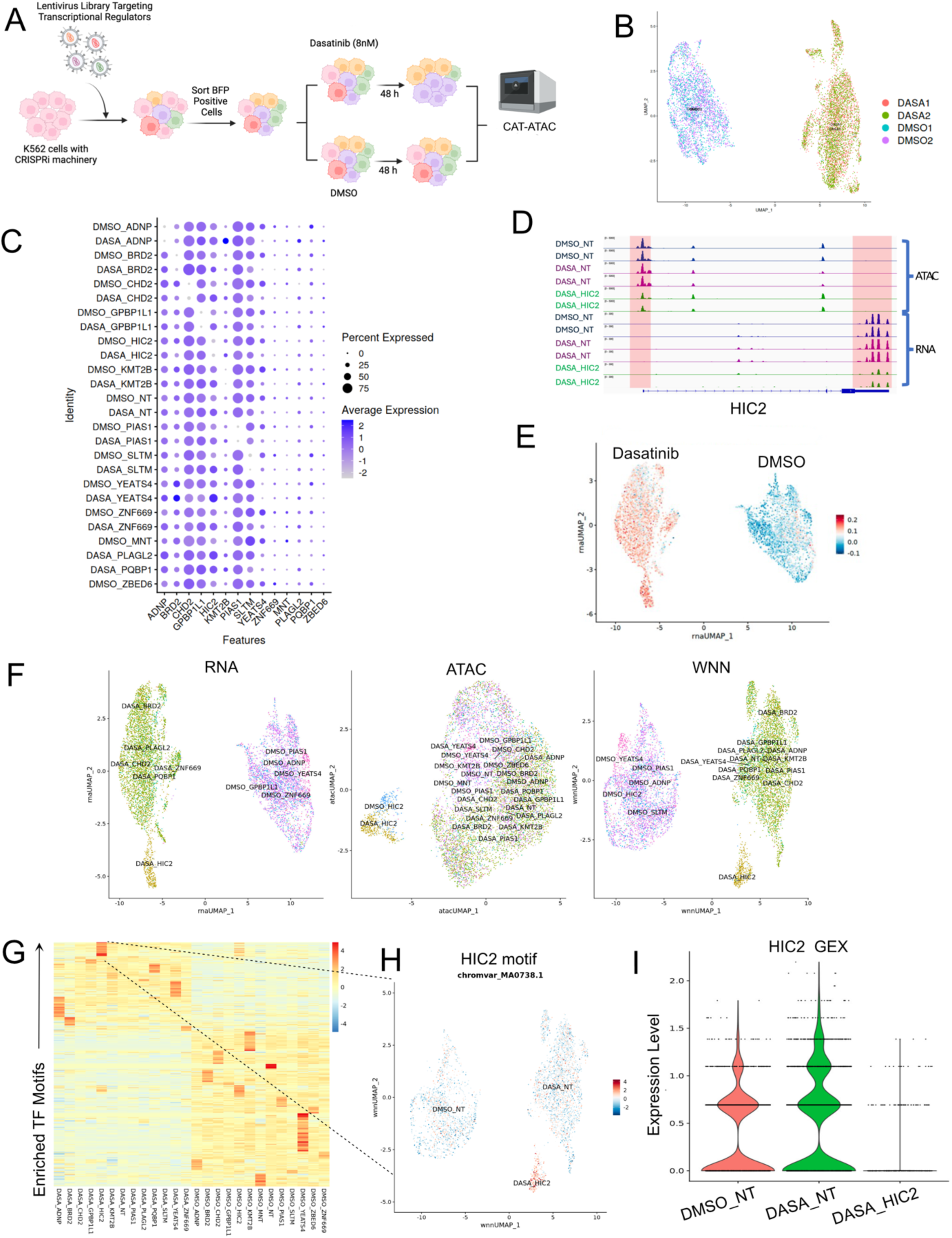
CAT-ATAC elucidates the combinatorial effect of gene and drug perturbation on RNA and ATAC profiles. A. Schematic showing the experimental outline for the CAT-ATAC experiment. B. UMAPs after sample integration showing replicate reproducibility. C. Dot plot showing target gene knockdown in cell populations which received the indicated guide in both dasatinib and DMSO treated conditions. D. Genome browser tracks for ATACseq and RNAseq peaks for DMSO_NT, DASA_NT and DASA_HIC2 cell groups at the HIC2 gene locus. Highlighted areas mark the promoter and 3’ end of the HIC2 gene. E. Module score plot showing that dasatinib treatment results in upregulation of erythrocyte progenitors genes identified by Velten et.al. F. UMAPs showing RNA, ATAC and weighted nearest neighbor profiles for cells expressing specific guideRNAs. G. Heatmap showing drug treatment and knockdown specific changes in chromvar motif activities. H. UMAP showing motif enrichment for HIC2 in the cell groups DMSO_NT, DASA_NT and DASA_HIC2. I. Violin plots showing RNA expression of HIC2 in the cell groups DMSO_NT, DASA_NT and DASA_HIC2.

Erythroid differentiation is one of the hallmarks of TKI treatment^35^. To assess whether dasatinib treatment in the experiment produced the expected effect on cells, we used the genes identified by Velten et al. 2017^36^ to assess erythroid differentiation in response to dasatinib treatment. We calculated a module score for these genes and saw an enrichment of the module in the dasatinib treated cells (Figure 3E) confirming the erythroid differentiation response and our ability to successfully assess transcriptional responses to dasatinib treatment in the context of genetic perturbations. We performed differential gene expression and chromatin accessibility analysis and observed perturbation-specific changes in gene expression and ATAC peaks (Supplemental Table 2, Supplemental Table 3, Supplemental Figure 4A and 4B). We observed that cells harboring knockdowns for the genes HIC2, KMT2B, YEATS4, PIAS1, PQBP1 and GPBP1L1 formed clusters when compared to the non-targeting guide expressing cells (Supplemental Figure 3C). Amongst these genes, HIC2 knockdown had the strongest effect on both the transcriptome and ATAC-seq profiles especially in the dasatinib-treated group (Figure 3F), leading us to focus on identifying genes downstream of HIC2 that might confer resistance to dasatinib. ChromVar^37^ can take sparse chromatin accessibility data from single cells as input and predict enrichment of transcription factor motifs. Using ChromVar^37^ we identified perturbation-specific enrichment of transcription factor motifs (Figure 3G). It has been previously reported that HIC2 has repressor activity^38^. Transcriptional repressors exhibit anticorrelation between motif activity and gene expression^39^, because chromatin bound by repressors is generally not accessible, and loss of repressor expression would expose more motifs normally occupied by the repressor. In agreement with these data, we find HIC2 motif to be enriched in cells where HIC2 expression is downregulated with CRISPRi (Figure 3H and 3I). Having the paired RNA-seq and ATAC-seq data from the same single cells, along with perturbation identities, was essential in establishing the anti-correlative relationship between HIC2 expression and motif enrichment. We thus demonstrate that, when applied in specific cellular contexts, CAT-ATAC can reveal novel context-dependent activation and repressive behaviors of transcription factors.

**Figure 4:**
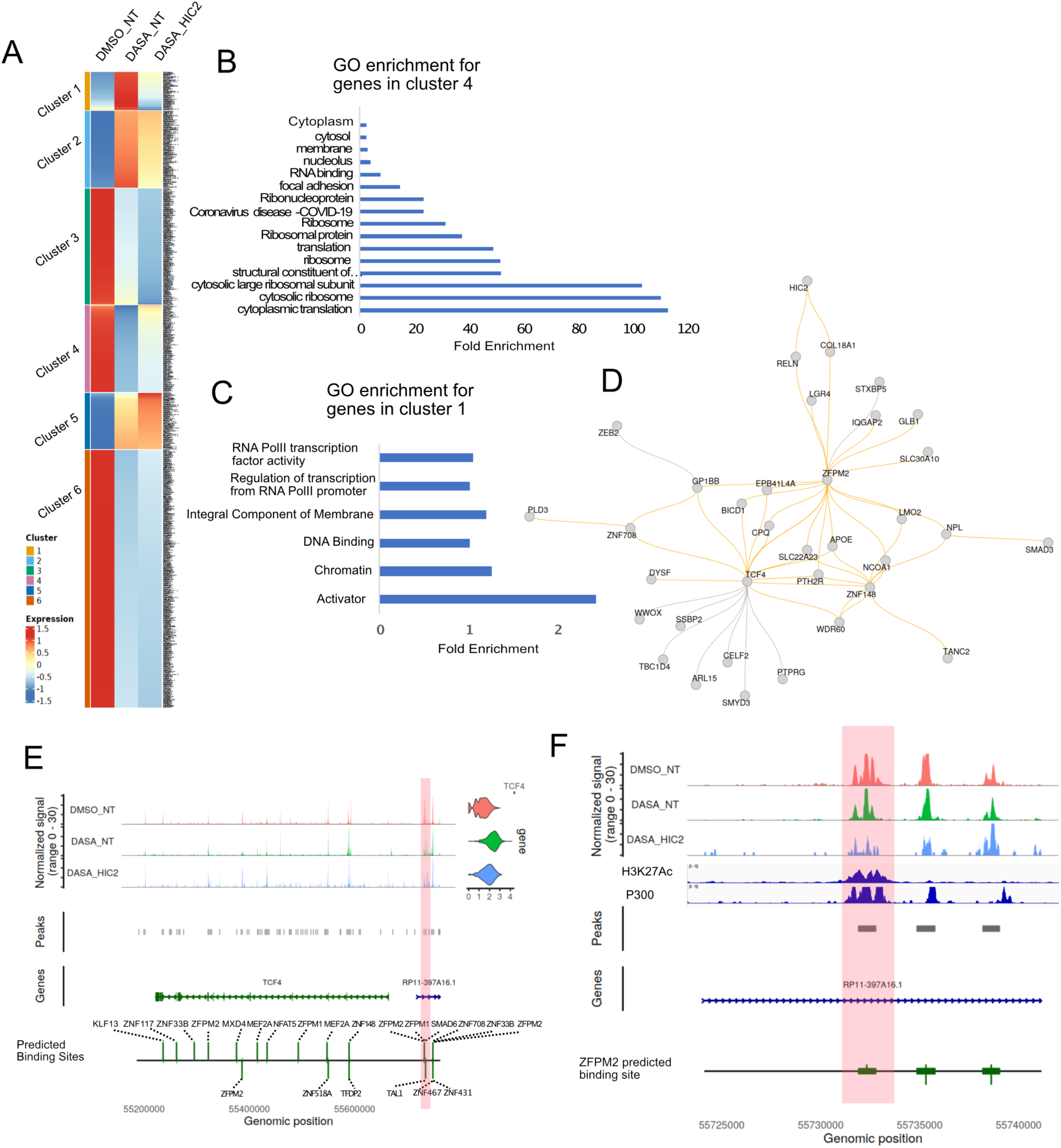
Identifying gene regulatory network underlying dasatinib resistance. A. Heatmap showing gene expression patterns in DMSO treated cells expressing non-targeting guides (DMSO_NT), dasatinib treated cells expressing non-targeting guides (DASA_NT) and dasatinib treated cells expressing HIC2 targeting guides (DASA_HIC2). B. GO analysis for genes in cluster 4 from panel A. C. GO analysis for genes cluster 1 from panel A. D. Subsetted gene regulatory network identified by querying the global network for genes differentially downregulated in dasatinib treated cells expressing HIC2 guides as compared to dasatinib treated cells expressing NT guides. E. Genome browser tracks showing the TCF4 locus. The first three tracks show ATACseq signal for the cell groups DMSO_NT, DASA_NT and DASA_HIC2, violin plots show the gene expression of TCF4 in the same cell groups. The last track shows the transcription factor binding sites predicted by Pando. Highlighted region shows ATAC-seq peaks upstream of TCF4 with predicted ZFPM2 binding sites. F. Zoomed in view of the highlighted region from panel E. The highlighted region in this track shows a ATACseq peak exhibiting reduced accessibility in DASA_HIC2 cell group. The next two tracks showing H3K27Ac and P300 enrichment in K562 cells were downloaded from ENCODE. The last track shows ZFPM2 binding sites as predicted by Pando.

### HIC2 indirectly activates genes potentially underlying dasatinib resistance

Since both our genome-wide CRISPRi screen and CAT-ATAC data identified HIC2 as a significant hit we went on to further identify genes controlled by HIC2 that drive dasatinib resistance. It was clear from the differential expression analysis that the HIC2 knockdown group had broadly altered gene expression patterns under dasatinib treatment conditions (Figure 3F), leading us to hypothesize that HIC2 targets’ expression are altered upon treatment with dasatinib. Considering that HIC2 downregulation sensitizes cells to dasatinib we hypothesized that some of the genes differentially expressed upon HIC2 knockdown should also affect sensitivity to dasatinib. Since HIC2 has repressor activity, the genes upregulated in the HIC2 knockdown cluster (DASA-HIC2 cells) would potentially be direct targets of HIC2. De-repression of these genes in K562 cells should sensitize them to dasatinib. In genes downregulated in DASA-HIC2 cells we expected that downregulation of these genes in K562 would sensitize the cells to dasatinib and their upregulation upon treatment with dasatinib could constitute the resistance response of the cells to dasatinib. We plotted the mean normalized read counts for the differentially expressed genes in the three cell groups, DMSO-treated cells expressing non-targeting control gRNAs (DMSO_NT), dasatinib-treated cells expressing non-targeting control gRNAs (DASA_NT) and dasatinib-treated cells expressing HIC2 gRNAs (DASA_HIC2) and clustered them using k-means clustering to identify patterns of gene expression changes (Figure 4A). We performed GO analysis for two sets of genes which follow Up-Down-Up (cluster 4) and Down-Up-Down (cluster 1) patterns of gene expression in the three conditions DMSO_NT, DASA_NT and DASA_HIC2 respectively (Figure 4B and 4C). We observed that genes following the Up-Down-Up pattern which potentially include direct targets of HIC2 are mostly involved with processes exhibiting post-transcriptional control of gene expression (Figure 4B) while genes following a Down-Up-Down pattern of gene expression, which may include dasatinib-resistance genes, were associated with transcriptional regulation functions (Figure 4C). As CAT-ATAC can only help us infer transcriptional control mechanisms we could not identify the genes controlled by direct targets of HIC2. Hence, we focused on identifying how the genes downregulated in the DASA-HIC2 cluster might contribute to dasatinib resistance via construction of gene regulatory networks.

### Gene Regulatory Networks identify novel pathways involved in dasatinib resistance

Recent single-cell GRN inference methods use paired scRNA-seq and scATAC-seq data to infer direct regulatory relationships between genes^40,41,42^. We used Pando^42^ to construct a gene regulatory network using scRNA and scATAC-seq data (Supplementary Figure 5). We queried this network for the genes downregulated in the DASA-HIC2 cluster and identified a sub-network comprising of 1^st^ degree connections to the genes in the query list (Figure 4D). The orange-colored connections show activating interactions while the grey connections represent repressive interactions. Upon inspecting the expression levels of genes in the subnetwork we found that majority of genes with activating interactions were upregulated in the DASA_NT cells as compared to DMSO_NT and DASA_HIC2 cells (Supplementary Figure 6A) while genes with repressive interactions did not follow this pattern of expression (Supplementary Figure 6B). We hypothesized that the genes with activating interactions in this sub-network are necessary for dasatinib resistance.

One such gene is ZFPM2, a transcription factor which is one of the central nodes in the sub-network in Figure 4D. The majority of interactions exhibited by ZFPM2 in this sub-network are activating interactions, leading us to hypothesize that ZFPM2 may be responsible for dasatinib resistance. One of ZFPM2’s activating interactors is the transcription factor TCF4. Upon inspecting the TCF4 locus we identified an ATAC peak upstream of TCF4 gene which was less pronounced in the DASA-HIC2 cells and harbored a predicted ZFPM2 binding site (Figure 4E and 4F). Decrease in accessibility of this peak correlated with a decrease in expression of the TCF4 gene in the DASA-HIC2 cell group (Figure 4E). We found that this peak overlaps with active histone marks H3K27Ac and P300 in K562 cells and is a potential active enhancer for TCF4 in K562 cells (Figure 4F). ZFPM2 could control TCF4 expression by binding an enhancer upstream of TCF4, an effector for the WNT signaling pathway^43^ which is known to be activated in the context of TKI resistance^44,45,46^. Thus, ZFPM2 may activate WNT downstream targets by controlling TCF4 expression, thereby conferring dasatinib resistance. Information about perturbation status of the target gene, ATAC-seq and RNA-seq data from the same cell helped us identify that HIC2 gene is necessary for ZFPM2 expression which in turn might contribute to activating TCF4 gene as a WNT pathway effector necessary for cell survival when treated with dasatinib. Paired RNA-seq and ATAC-seq data from CAT-ATAC allowed for the identification of the cis-regulatory relationship between the TCF4 promoter and accessible DNA site upstream of TCF4. CAT-ATAC thus provides the ability to build tesTable mechanistic hypothesis by combining multiple data types from different aspects of cellular regulatory processes.

### ZFPM2 is necessary for increased dasatinib resistance

To determine whether ZFPM2 is necessary for dasatinib resistance, we knocked down HIC2 (positive control), and ZFPM2 using CRISPRi in K562 cells (guide sequences in Supplemental Table 1). qRT-PCR analysis showed significant knockdown of gene expression for both genes when compared to non-targeting controls (Figure 5A). We next performed flow cytometry analysis post-dasatinib treatment to elucidate the effect of knockdowns on dasatinib sensitivity. The gating strategy for the flow cytometry analysis is illustrated in Supplemental Figure 7. After treating the cells with 8nM dasatinib for 72 hours, we observed that the cells with gene knockdowns showed a significant decrease in survival compared to the non-targeting control expressing cells thereby indicating these genes caused increased resistance to dasatinib (Figure 5B). We also performed a longitudinal timecourse experiment quantifying % viability of cells over time and saw significant decrease in viability for DASA-HIC2 and DASA-ZFPM2 cells across all the timepoints tested (Figure 5C). Thus, we can conclude that ZFPM2 confers dasatinib resistance.

**Figure 5:**
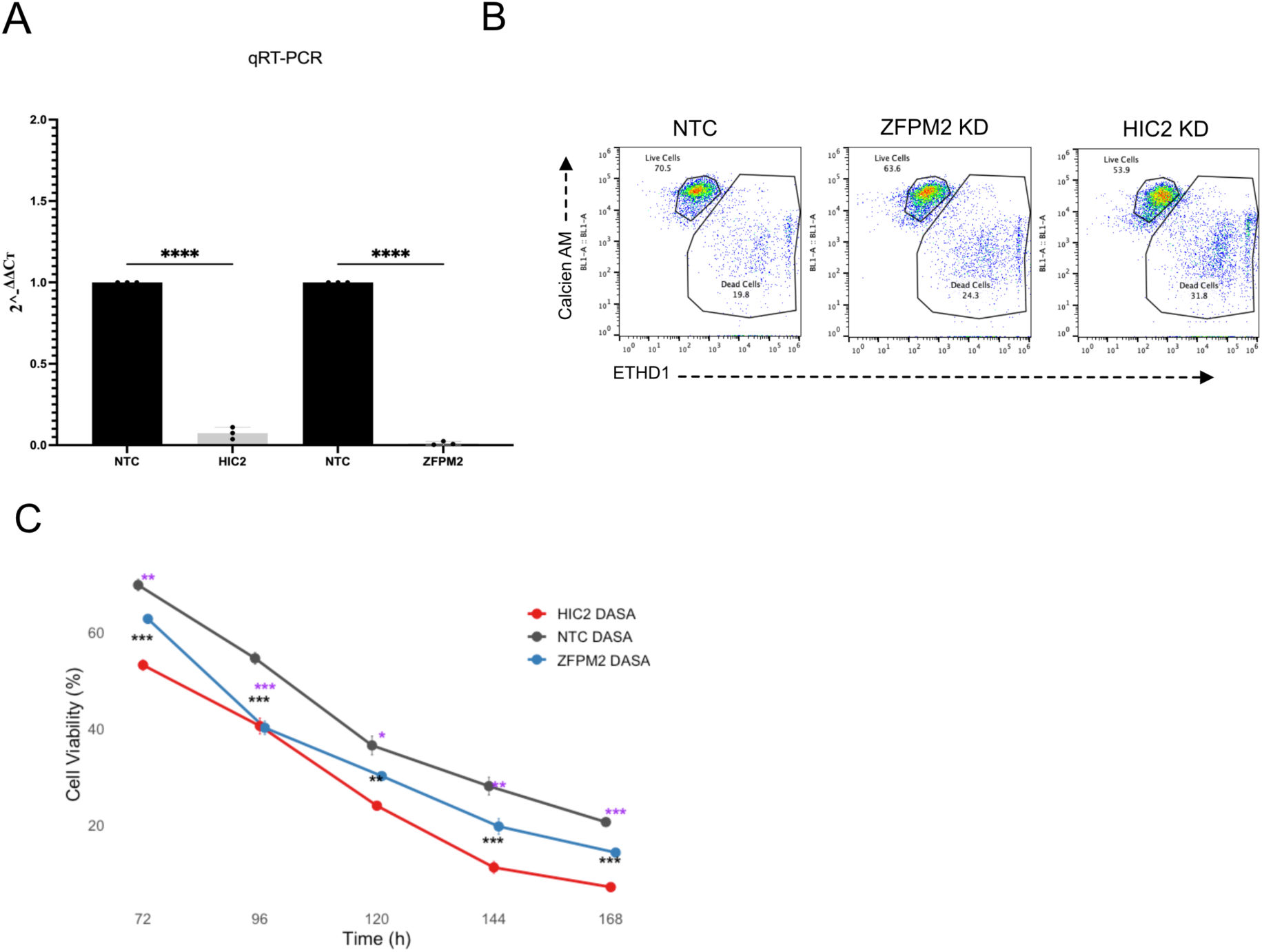
ZFPM2 and PTH2R are necessary and sufficient for increased resistance to dasatinib. A. qPCR data showing CRISPRi mediated knockdown of gene expression for the genes HIC2, PTH2R and ZFPM2. B. Flow-cytometry plots at 72h post dasatinib treatment showing the distribution of cells for each of the three gene knockdown conditions along with control cells treated with dasatinib and stained with Calcien-AM and EthD-1 to identify live and dead cells respectively. C. Timecourse experiment showing flow-cytometry data from 72h to 168h post dasatinib treatment quantifying % viable cells in HIC2 KD, ZFPM2 KD and control cells. Each data point represents the mean ± SD of three biological replicates (n=3). Statistical significance was assessed at each time point using unpaired two-tailed t-tests comparing HIC2 KD or ZFPM2 KD vs. NTC. Black asterisks indicate significant differences for HIC2 KD vs. NTC, while purple asterisks indicate significant differences for ZFPM2 KD vs. NTC (* corresponds to p < 0.05, ** corresponds to p < 0.01 and *** corresponds to p < 0.001).

## Discussion

We have shown that CAT-ATAC robustly captures guide RNAs, resulting in up to 77% of guide assignment in tested cell lines, while producing high-quality scRNA- and scATAC-seq data. The main application of this approach is to establish a novel platform for conducting CRISPR screens, by combining small-scale to genome-wide gene perturbations with massively parallel sequencing of RNA and chromatin accessibility at the single-cell level. CAT-ATAC generates paired gene expression and open chromatin data without the need to perform separate pooled screens for each of the modalities, thus reducing the avenues for technical and biological confounders. The experiments in this study used a primer containing the CS1 direct capture sequence to reverse transcribe guide RNAs (Figure 1B), but this assay is designed to be universally compatible with any type of CRISPR perturbation or synthetic construct by including a splinted reverse transcription primer against the targeted sequence. Two recent studies have demonstrated alternative approaches to perform CRISPR screens with multimodal gene expression and open chromatin readouts^47,48^. However, these methods would either require re-cloning most labs’ existing CRISPR guide libraries into a modified vector^47^ or implementing complicated experimental approaches with multiple rounds of barcoding^48^. CAT-ATAC offers a straightforward and easy-to-perform workflow, incorporating minimal modifications to the 10X Genomics Multiome assay and is compatible with existing designs of the CRISPR gRNA libraries while reporting comparable capture efficiencies of up to 77%, which could likely be optimized further by employing enrichment strategies using biotinylated PCR primers similar to that used for MultiPerturb-seq^48^. Additionally, the ability to directly capture guide RNAs enables the use of constructs that express multiple guide RNAs targeting one or several genes^23,49,50^ which makes the studies of combinatorial perturbations feasible. Our chemical-genetic screen identified genetic modifiers that sensitize cells to dasatinib. These genetic modifiers included several genes with transcriptional regulatory activity. We show for the first time that HIC2 expression drives cellular resistance to dasatinib. Our results support an observation from a study comparing the gene expression profiles of CD34+ cells from CML patients and controls which reported overexpression of HIC2 in patient cells suggesting a role for HIC2 in disease progression^51^. When applied to identifying cancer drug resistance mechanisms CAT-ATAC was able to identify a gene regulatory network that is activated under dasatinib treatment and requires expression of HIC2 gene, and to further identify ZFPM2 as a central transcription factor in this network. Using the scATAC and scRNA-seq data from HIC2 knockdown cells we were able to predict a novel mechanism whereby ZFPM2 putatively binds a cis-regulatory element upstream of TCF4, thereby controlling expression of TCF4, a transcription factor known to be an effector for the WNT signaling pathway that contributes to TKI resistance for CML and other myeloid leukemias.

Besides capturing guide RNAs in CRISPR screens, this method is further extendable to any other nucleic acid target, such as a reporter DNA or RNA, BCR/TCR immunoglobulin sequences, or DNA conjugated to antibodies and antigens which can be used to measure protein abundance along with gene expression and chromatin accessibility. These types of multiparametric single-cell readouts will be transformative to shape our understanding of fundamental biological processes, cell fate decisions and disease mechanisms by harnessing the heterogeneity inherent to complex cell populations instead of reducing datasets to bulk analysis. By providing a better understanding of the link between DNA accessibility, gene transcription, and protein translation and processing, these approaches can be broadly applied to probe regulation by non-coding regions of the genome, allowing for more systematic studies of the links between gene regulatory elements and gene expression. In addition, these tools can be leveraged to gain biological and mechanistic insights into the relationship between recognition motifs, transcription factor binding, and downstream gene activity, enabling further exploration of the dynamics of transcription factor-driven cell fate changes. Furthermore, these types of multimodal perturbation datasets will prove invaluable for building foundational models in biology such as the virtual cell^52^.

## Data availability

Raw and processed CAT-ATAC sequencing data can be downloaded from Gene Expression Omnibus (<u>GSE288996</u>). Data from previously published studies were from the Sequence Read Archive or Gene Expression Omnibus: SHARE-seq^53^ (<u>GSE140203</u>), SNARE-seq^54^ (<u>GSE126074</u>), Paired-seq^55^ (<u>GSE130399</u>), sci-CAR-seq^56^ (<u>GSE117089</u>) and MultiPerturb-seq^48^ (<u>GSE277747</u>). The human genome hg38 (GENCODE v32/Ensembl98) was from 10x Genomics (https://cf.10xgenomics.com/supp/cell-exp/refdata-gex-GRCh38-2020-A.tar.gz).

## Code availability

All the code generated for analysis of the CAT-ATAC data including for counting guides, guide to cell assignment and downstream differential expression and gene regulatory analysis will be made available upon request to the corresponding authors and released upon publication of this manuscript.

## Acknowledgements

We thank the UCSF Center for Advanced Technology (CAT) for NGS sequencing services. Sequencing was performed at the UCSF CAT, supported by UCSF PBBR, RRP IMIA, and NIH 1S10OD028511-01 grants. We would also like to thank LGR team members Greyson Lewis, Sailaja Peddada, and Adam Litterman for their advice and Lauren Enriquez for help in setting up automation protocols for guide cloning. Some Figure elements were created using BioRender.com.

## Funding

This work is supported by the Laboratory for Genomics Research established by GSK, UCSF, and UC Berkeley.

## Author Contributions

K.S, Y.A.Y, E.D.C and L.P contributed to the study’s overall conception, design, interpretation and co-wrote the manuscript. E.D.C conceptualized CAT-ATAC, Y.A.Y performed the CAT-ATAC optimization and method development experiments. K.S designed the screening concept. K.S and K.M worked on genome-wide dasatinib screen and CAT-ATAC follow-up screen. Y.A.Y, K.S. and K.F performed data analysis and interpretation. V.S, K.F and S.F wrote the data analysis pipeline. J.P wrote the code for guide RNA assignment. G.A helped in making the multiome libraries. K.M and H.E worked on validation experiments. R.M cloned arrayed guide plasmids. R.E, K.S, E.D.C and S.S helped Y.A.Y troubleshoot CAT-ATAC method development experiments.

## Conflict of Interest

Y.A.Y, V.S, J.P, R.E, G.A and S.S are employees of GSK. E.D.C. is a co-founder of Survey Genomics.

## Methods

### Cell lines

The human biological samples were sourced ethically, and their research use was in accord with the terms of the informed consent under an IRB/EC approved protocol. iPSC cell line (Proprietary for GSK use generated by Takara) was engineered to express doxycycline inducible CRISPRa machinery (dCas9-VP64-p65-Rta). The CRISPRa machinery was integrated into the AAVS1 safe harbor locus to minimize silencing.

The K562-CRISPRi cell line was a gift from the Gilbert lab. K562-CRISPRi line was maintained in the standard conditions of DMEM+10% FBS with antibiotics as recommended by ATCC.

### gRNA cloning

For the pilot experiment top two guide RNAs from the Weissman V2 CRISPRi library targeting the 3 transcription factors (GATA1, MEOX1 and NGN2) and were cloned in dual guide lentiviral construct compatible with Perturb-seq using Gibson assembly. For the arrayed validation of hits from the dasatinib screen top two guides which exhibited the highest phenotype score for sensitizing the cells to dasatinib were cloned individually in the dual guide gRNA vector using Gibson assembly. Gibson assembly process was as follows:

Dual guide RNAs were cloned in a lentiviral vector backbone using Gibson assembly cloning. An insert DNA sequence with flanking BsmBI restriction sites and containing a constant region with Perturb-seq compatible capture sequence 1 (CS1: GCTTTAAGGCCGGTCCTAGCAA) was cloned into an intermediate transfer vector and transformed into bacteria for plasmid amplification (See insert sequence below). To prepare a linear insert for cloning the amplified plasmid was digested using BsmBI restriction digestion and the desired fragment was purified using gel purification. The backbone vector was also linearized using BsmBI restriction digestion. This backbone vector (pLGR134) was derived from pLGR002 (Addgene #188320) by swapping out the BstXI and BlpI sites for a pair for BsmBI sites. ssDNA oligos for guides were ordered with 20 base pair overhangs on both 5’ and 3’ sides which partially overlap the constant region and mU6 promoter for guide 1 and constant region and hU6 promoter for guide 2. Linearized vector, insert and guide oligos were then assembled into dual guide vectors in a Gibson assembly reaction. Colonies were validated by Sanger sequencing.

Insert sequence before BsmBI restriction digest: CGTCTCAagaggtttcAGAGCTAAGCACAAGAGTGCATAGCAAGTTGAAATAAGGCTAGTCCGTTTACAACTTGGCCGCTTTAAGGCCGGTCCTAGCAAGGCCAAGTGGCACCCGAGTCGGGTGCTTTTTTTGCTCGAATCTACACTCAGCTATGGCGCGCCCCAAGGTCGGGCAGGAAGAGGGCCTATTTCCCATGATTCCTTCATATTTGCATATACGATACAAGGCTGTTAGAGAGATAATTGGAATTAATTTGACTGTAAACACAAAGATATTAGTACAAAATACGTGACGTAGAAAGTAATAATTTCTTGGGTAGTTTGCAGTTTTAAAATTATGTTTTAAAATGGACTATCATATGCTTACCGTAACTTGAAAGTATTTCGATTTCTTGGCTTTATATATCTTGTGGAAAGCCAgaaacatgGAAAGGAGACG

Plasmids for the vectors targeting the 20 selected genes along with 6 NTCs required for the pooled multiome screen were then mixed in equimolar amounts to generate a pooled plasmid library.

### Lentivirus production and transduction

For all arrayed experiments lentivirus was generated in an arrayed format by transfecting LentiX cells (Takara 632180) with individual gRNA vectors. To generate the lentivirus for transducing iPSC cells LentiX cells were forward transfected with the gRNA vectors and lentiviral packaging plasmids (dR8.91 and MD2G) using the Mirus transfection reagent (MIR 2700). The three plasmids were mixed in 1:1:0.1 ratios for gRNA vector, MD2G and dR8.91 respectively. The lentivirus was then harvested 48hr post transfection in MTeSR1 medium for transduction in iPSCs. The lentivirus for arrayed validation experiment was generated in a 96 well format where the LentiX cells were reverse transfected using TransIT-VirusGEN® Reagent (MIR 6704). The gRNA and packaging plasmids were mixed in the same 1:1:0.1 ratio followed by lentivirus harvest in media with 10% FBS. Pooled lentiviruses were generated in either 6 well or 15 cm dishes depending on the amount of lentivirus required. LentiX cells were forward transduced using Mirus transfection reagent and the plasmids were mixed in the ratios mentioned above. Lentivirus was then harvested in media containing 10% FBS.

### Genome-wide CRISPRi screen

K562 cells containing dCas9-KRAB CRISPRi machinery were infected with CRISPRi V2 library containing the top5 guides (Addgene 83969) to achieve 1000-fold coverage at an MOI of 0.3. Transduced cells were selected with puromycin until 90% of the cell population showed BFP expression indicating guide expression. Cells were then treated with a single dose of 8nM dasatinib (determined from a drug titration experiment) for 72hr. This was followed by recovery for 5 days. An equivalent number of transduced cells were treated with DMSO as a vehicle control. Both the dasatinib and DMSO treated cells were then collected and genomic DNA was isolated using the Macherey-Nagel Nucleospin Blood XL kit (740950). The cassette encoding the gRNAs was then amplified by PCR and gRNA abundance was determined by next generation sequencing (NGS) as previously described^57,58,59^. The libraries were sequenced on an Illumina NextSeq550 sequencer and the sequencing data was analyzed using the screen processing pipeline^57^.

### Arrayed validation of the hits from CRISPRi screen

A dual-guide lentiviral library targeting 60 genes and two non-targeting controls was prepared in-house as described above and used for arrayed lentiviral generation in HEK 293T cells. K562-CRISPRi cells were then “spinfected” at 1000 RCF for 1 hour at 22C with this lentivirus in an arrayed fashion in the presence of 1 ug/mL of polybrene, left overnight at 37C, and then selected via puromycin at a concentration of 2ug/mL for 48 hours. Following selection, guide-infected cell lines were treated with 8nM Dasatinib and their viability was recorded using Propidium Iodide nuclear stain (Invitrogen P1304MP) every 24 hours following treatment on an Invitrogen Attune NxT Flow Cytometer.

### CAT-ATAC

#### CAT-ATAC pilot experiment in iPSCs

iPSCs were transduced at MOI of 0.3 in the presence of 1 ug/mL of polybrene overnight in an arrayed format, followed by puromycin selection for one week at a concentration of 2 ug/mL. Three transcription factors were overexpressed (NEUROG2, GATA5, MEOX1) by supplementing doxycycline (4uM) in the media (mTeSR1, StemCell Technologies) for 7 days prior to CAT-ATAC analysis. As a control, a non-targeting scrambled gRNA sequence was used. After 7 days of overexpression cells were dissociated into a single-cell suspension using Accutase (ThermoFisher Scientific A1110501), counted and then equal numbers of cells per perturbation were mixed to make a pool of cells prior to processing for CAT-ATAC.

#### Pooled Lentivirus Generation and Dasatinib Treatment

A pooled lentiviral library containing 26 dual-guide vectors targeting 20 previously identified transciptional regulatory genes and 6 non-targeting genes was generated as described previously. The resulting virus was then titered to determine the amount of virus required for desired MOI of 0.1 in K562 cells. Accordingly, 1.5*10^7 cells containing CRISPRi machinery were spun down and resuspended directly into 2.4mL of this pooled virus and “spinfected” at 1000 RCF for 1 hour at 22C in the presence of 1 ug/mL of polybrene and incubating overnight at 37C. Guide infected cells were sorted via FACS based on BFP expression and allowed to recover for 5 days. 2.5*10^7 of these sorted cells were then aliquoted into 4 shaking flasks, two of which were treated with Dasatinib and two of which were treated with DMSO for 48 hours before being collected and aliquoted for CAT-ATAC.

#### CAT-ATAC method

Detailed CAT-ATAC protocol is in Supplemental Text 1. Briefly cells were harvested, nuclei were extracted and tagmented according to the 10X Multiome protocol. During the GEM Generation step, a duplex splint oligo is spiked into the barcoding reaction mix to allow reverse transcription of gRNAs and ligation onto the capture oligo of 10X Multiome gel beads. The resulting cDNA from captured guide RNAs is amplified to make an Illumina-compatible sequencing library containing P5 and P7 adaptors as well as sample indexes. GEX libraries were sequenced on NovaSeqX with paired-end sequencing, 28×10×10×90, at a minimum of 20,000 read pairs/cell. ATAC and CRISPR libraries were pooled at a 5:1 ratio and were sequenced on NovaSeqX with paired-end sequencing, 151×8×24×151 (since at least 75bp PE is needed to be able to sequence through protospacer region for CRISPR libraries), at a minimum of 25,000 read pairs/cell for ATAC libraries and 5,000 read pairs/cell for CRISPR libraries.

### CAT-ATAC data analysis

#### Processing FASTQ files after NGS

We extracted the protospacer-cell barcode pairs from the guide capture FASTQ files using a custom Bash script that first finds the reads containing both the protospacer and capture sequence via BBDuk alignment (https://sourceforge.net/projects/bbmap/) at a Hamming distance=1. The matching reads are then demultiplexing based on unique UMIs using a guide calling algorithm described in the Guide Assignment section, and count tables are generated. GEX and ATAC FASTQs were aligned to GRCh38 version of the human genome using prebuilt packages for Cell Ranger ARC v.2.0.1. Post alignment filtering, barcode counting and counting of GEX and ATAC molecules in the FASTQ files was performed using the cellranger-arc count function.

#### Guide Assignment

For the pilot CRISPRa experiment in iPSCs cells were assigned a guide identity using HTOdemux function in Seurat which is based on a previously published algorithm that was developed for cell hashing oligos^60^. We found that this method had low signal to noise ratio in our pooled CRISPRi screen, so we developed a new algorithm to assign guides to cells. For the pooled CRISPRi CAT-ATAC screen the UMIs per cell for each guide are modeled using a mixture of a zero-inflated negative binomial distribution and a negative binomial distribution (without zero-inflation). To maintain identifiability, the parameter space is constrained so that the negative binomial distribution without zero inflation has a larger mean than the negative binomial component of the zero-inflated distribution. Estimates for the negative binomial parameters and mixture proportions were obtained by maximum likelihood and used to determine the posterior probability that each observation arose from the negative binomial component with the larger mean. A cell was classified as expressing a guide if the posterior probability that the corresponding UMI measurement came from the negative binomial component with the larger mean was greater than 0.5.

#### Data QC, filtering, integration and differential testing

Seurat v.4.3.0^29^ was used for subsequent analysis. Cells with mitochondrial RNA percentage > 20%, RNA count <1000 and >40000, nucleosome signal > 2, and TSS enrichment <1 were filtered out. Seurat SCTransform v2.0 was used to normalize and scale the data as well as to find variable features. Cell cycle-associated genes were also regressed out using Seurat’s CellCycleScoring SCTransform functions to mitigate their effects on downstream analyses. We then ran Mixscape on each sample to calculate perturbation-specific signatures for each cell. This step included the identification and exclusion of cells that evaded CRISPR-mediated perturbation. The method also facilitated the visualization of similarities and differences among cells subjected to various perturbations.

The top 3,000 highly variable genes were used for PCA and UMAP dimensionality reduction. Libraries were integrated using Seurat’s merge and IntegrateEmbeddings functions. Cells were clustered using the FindClusters function and the Louvain algorithm. To identify significant changes associated with each perturbation, differential expression testing was conducted using Wilcoxon rank sum test through Seurat FindMarkers function and differential accessibility testing was conducted using logistic regression in Signac.

#### Module score analysis

Genes associated with erythroid differentiation were identified from Velten et.al. ^36^. Module scores for these sets of genes were calculated using AddModuleScore function in Seurat. These module scores were then plotted on the rnaUMAP to visualize the enrichment of the module in specific treatment groups.

#### H3K27Ac and P300 ChIPseq data

Bigwig files for H3K27Ac (ENCSR000AKP) and P300 (ENCSR000EGE) ChIPseq datasets in K562 were downloaded from ENCODE (The Encyclopedia of DNA Elements) consortium.

#### Gene Ontology analysis

Selected gene sets either identified as hits in the CRISPR screens or as significant differentially expressed genes were used for gene ontology analysis as mentioned in the results section. Gene ontology analysis was performed using the DAVID knowledgebase (https://davidbioinformatics.nih.gov/).

#### Motif enrichment and footprinting analysis

Motif and footprinting analysis were performed using Signac’s AddMotifs, FindMotifs, and FootPrint functions. Motif activities were calculated using Signac’s RunChromVAR function and JASPAR2024 database^61^.

#### Gene Regulatory Network inference with Pando

To perform gene regulatory network inference, we used the functions initiate_grn, find_motifs, infer_grn, find_modules, and NetworkModules from the Pando package^42^ . We first built a global GRN including all the cells in the integrated multiome object. To identify HIC2 specific network we subset this global GRN using a list of genes that were differentially downregulated in DASA_HIC2 cells as compared to DASA_NT cells.

#### Validation of ZFPM2

The top two guide RNAs from the Weissman V2 CRISPRi library targeting ZFPM2 gene were cloned into the LGR dual guide vector as described earlier. Lentivirus was generated as described before and concentrated using Lenti-X™ Concentrator (cat # 631232). K562 cells containing CRISPRi machinery were then transduced with guide lentiviruses for HIC2 (positive control), a non-targeting control (NTC) and ZFPM2. The transduced cells were selected with puromycin and then used for downstream validation experiments.

#### Validation of target knockdown post CRISPRi

Total RNA was extracted from K562 CRISPRi constructs targeted against NTC, HIC-2, and ZFPM2 (*n* = 3/each group) by using RNeasy Plus Mini Kit (Cat # 74134) and converted to cDNA using SuperScript Vilo cDNA synthesis kit (Cat # 11754050). A total of 1 μg RNA was converted to cDNA. cDNA was amplified using a SYBR-green qPCR master mix (Cat #43-676-59) and specific primers in Supplemental Table 4. GAPDH was used as the reference gene.

The PCR steps consisted of an initial activation 50°C for 2 mins, pre-soak step at 95 °C for 10 min, denaturation of at 95 °C for 15 sec, followed by annealing step at 60 °C for 1 min for 40 cycles. The last step is the melting curve (95°C for 15 sec, 60°C for 15 sec, and 95°C for 15 sec). Detection, quantification, and data analysis were performed in the CFX manager real-time detection system (Bio-Rad). The Ct values were calculated for each gene and were compared with the reference gene GAPDH, followed by estimation of ΔΔCt and fold change (2^−ΔΔCt^) to assess the relative gene expression.

#### Dasatinib treatment and cell viability analysis

Dasatinib (Millipore Sigma, cat. # SML2589) was diluted with DMSO to final concentration of 8nM, 0.01% DMSO was used as a vehicle control. K562 cells expressing either NTC, HIC2 or ZFPM2 guides were treated with 8nM dasatinib for 72 hr. Live-dead staining was performed using 2uM calcien-AM (cat # C1430) and 2.5uM ETHD1 EthD-1 (cat # E1169) and cells were analyzed using the Attune NextGen flow cytometer (Thermo Scientific, USA) at 72h, 96h, 120h, 144h and 168h post dasatinib treatment. FSC and SSC dot plot was used to gate out the debris present in the suspension and percentages of live (calcien-AM positive) and dead (EthD-1 positive) cells were estimated.

## Supplementary Figures

**Supplementary Figure 1:**
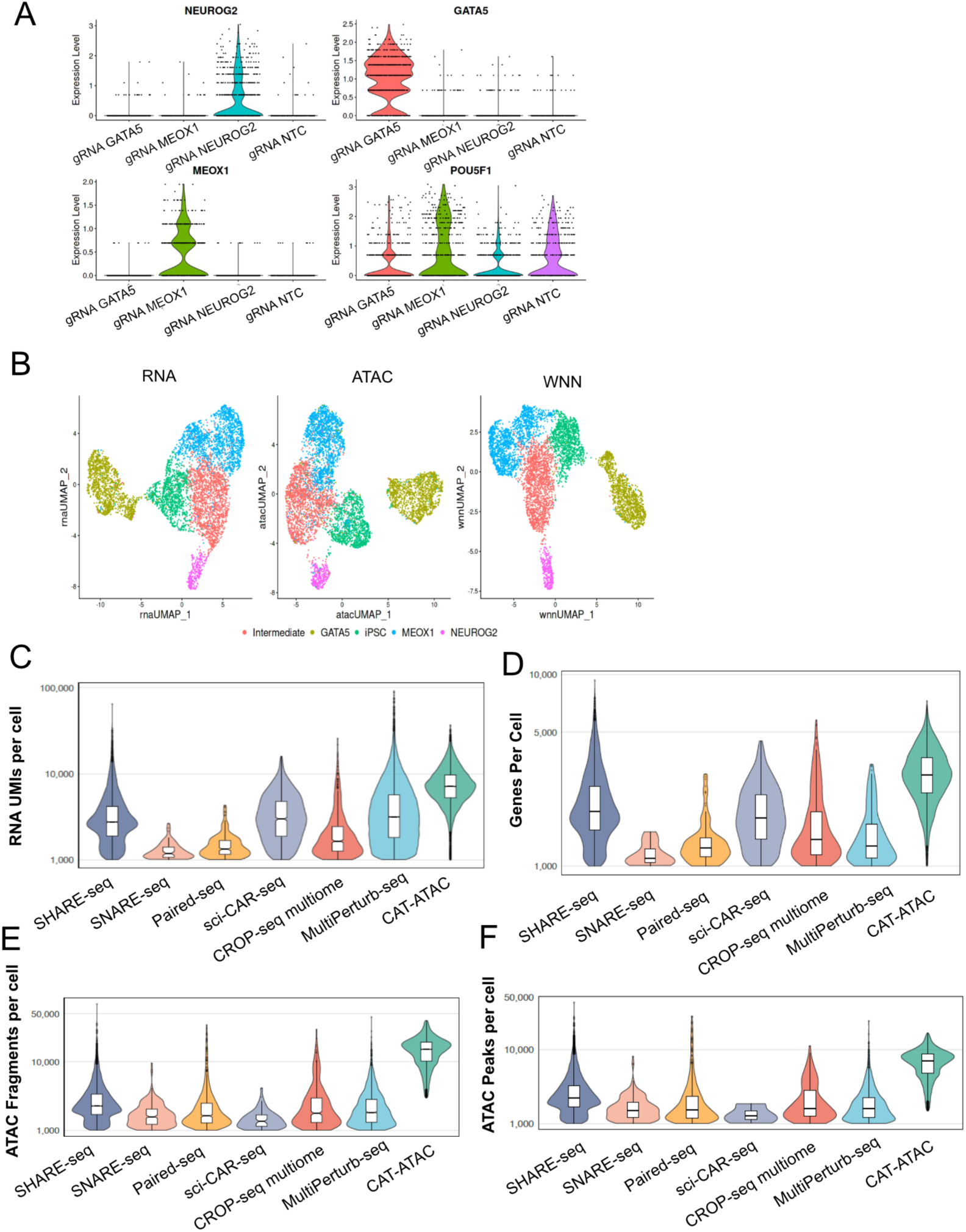
CAT-ATAC QC and comparisons with other technologies A. Violin plots showing protospacer expression in clusters shown in UMAP in Figure 1 panel F. B. UMAPs based on RNA, ATAC and joint profiles for CAT-ATAC in iPSCs with 3 TFs (GATA5, NEUROG2, MEOX1) and 1 non-targeting control overexpressed by CRISPRa. C. Violin plots comparing RNA UMIs per cell metric for various Multiome approaches with CAT-ATAC. D. Violin plots comparing Genes per cell metric for various Multiome approaches with CAT-ATAC. E. Violin plots comparing ATAC fragments per cell metric for various Multiome approaches with CAT-ATAC. F. Violin plots comparing ATAC peaks per cell metric for various Multiome approaches with CAT-ATAC.

**Supplementary Figure 2:**
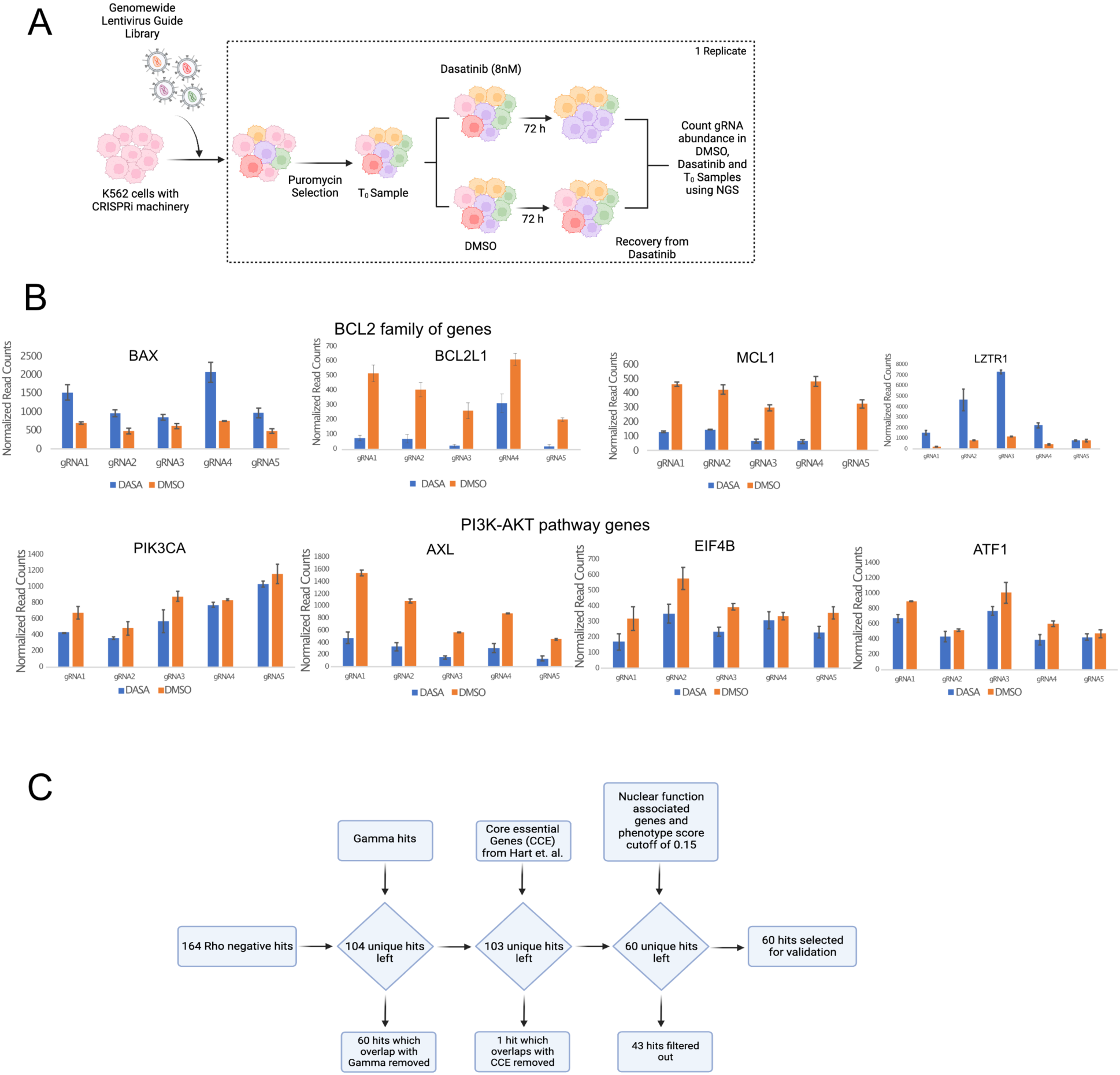
Dasatinib genomewide screen design and QC A. Design of the dasatinib survival screen. B. Normalized read counts of guides targeting some of the genes that were identified as hits and were highlighted in Figure 2 panel B. Error bars represent standard deviation. C. Strategy to filter the sensitizing hits.

**Supplementary Figure 3:**
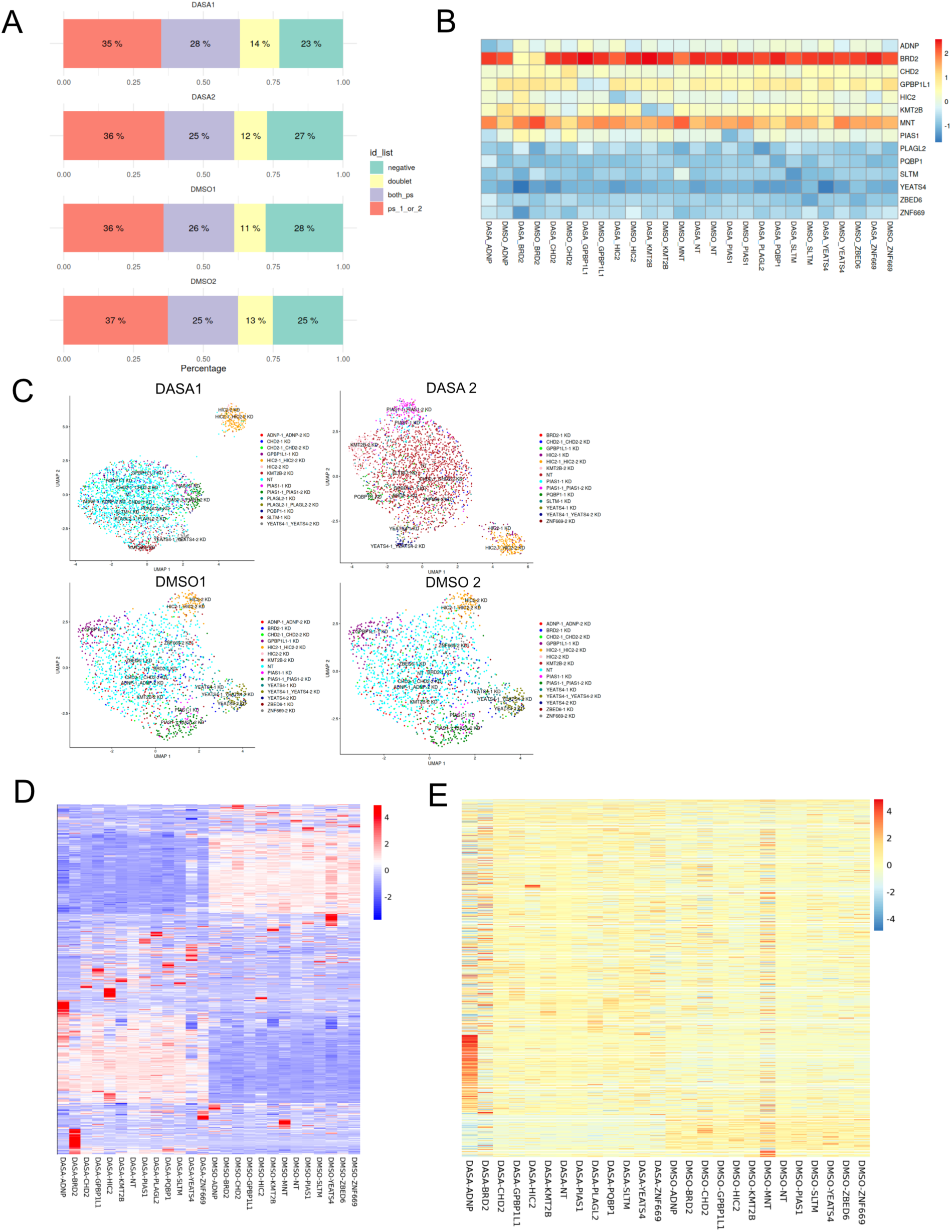
CAT-ATAC QC for dasatinib screen A. Bar graph showing guide assignment for each of the 4 samples from the 10x Multiome screen. B. Heatmap showing average accessibility at the promoters of targeted genes in both dasatinib and dmso treated samples. C. UMAPs showing RNA expression profiles of cells for individual samples grouped by perturbation classes based on gRNA expression. D. Heatmap showing top 1000 differentially expressed genes from each of the perturbation classes across the two treatment groups dasatinib and DMSO. E. Heatmap showing differentially accessible ATAC peaks from each of the perturbation classes across the two treatment groups dasatinib and DMSO.

**Supplementary Figure 4:**
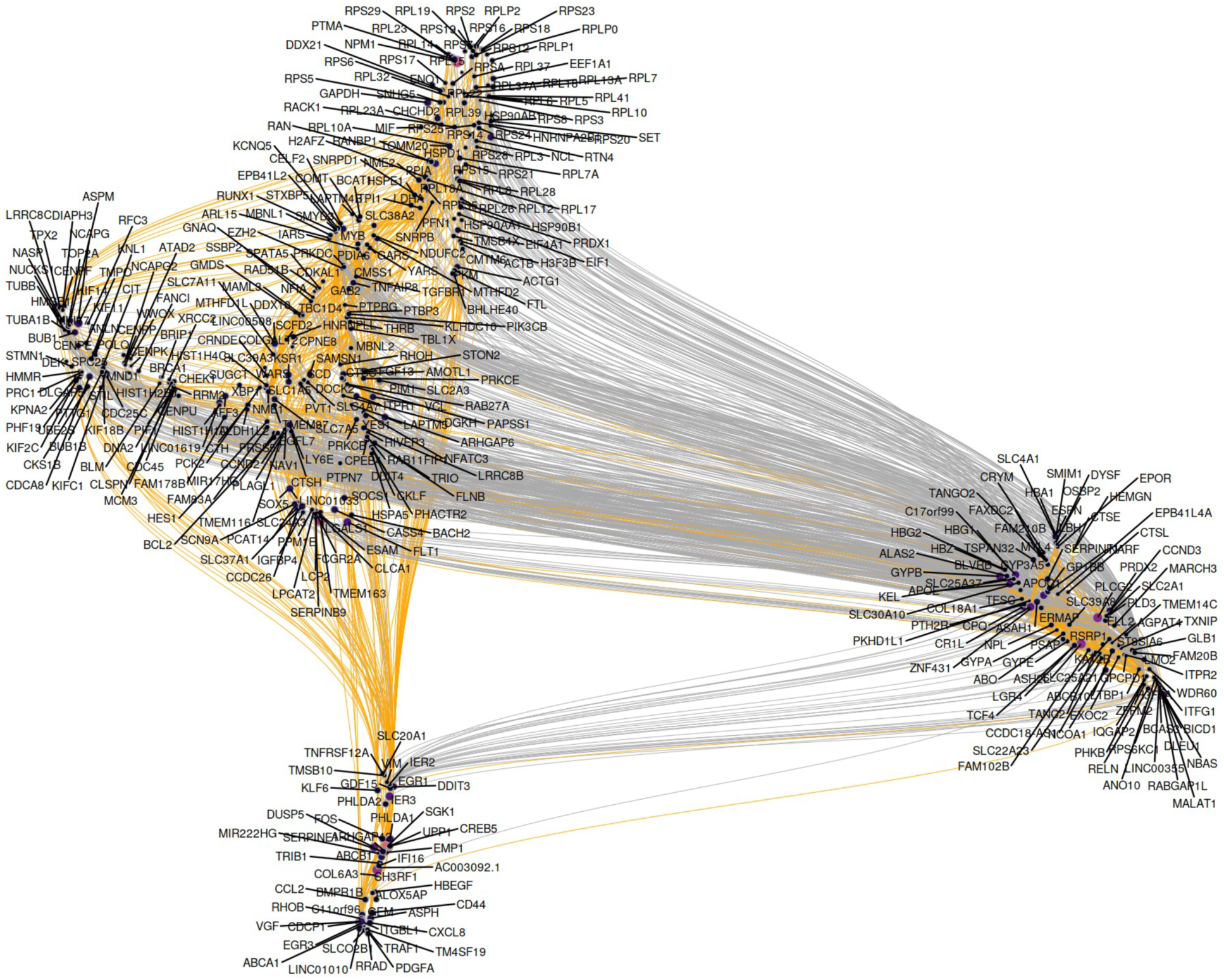
Global GRN Global GRN constructed using Pando

**Supplementary Figure 5:**
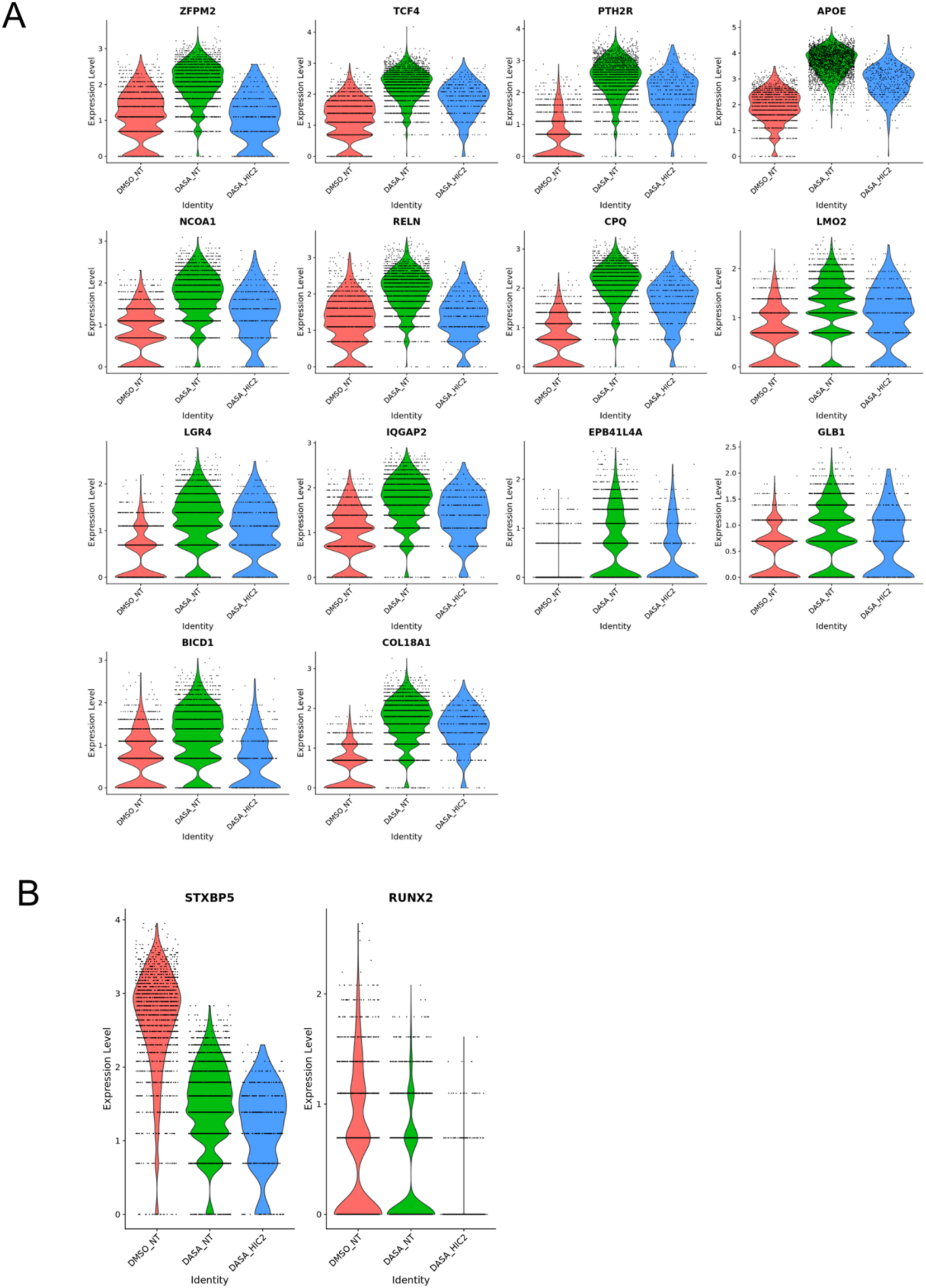
Expression of genes in the subsetted GRN A. Violin plots showing expression of genes having activating connections in the subsetted gene regulatory network. B. Violin plots showing expression of genes having repressive connections in the subsetted gene regulatory network.

**Supplementary Figure 6:**
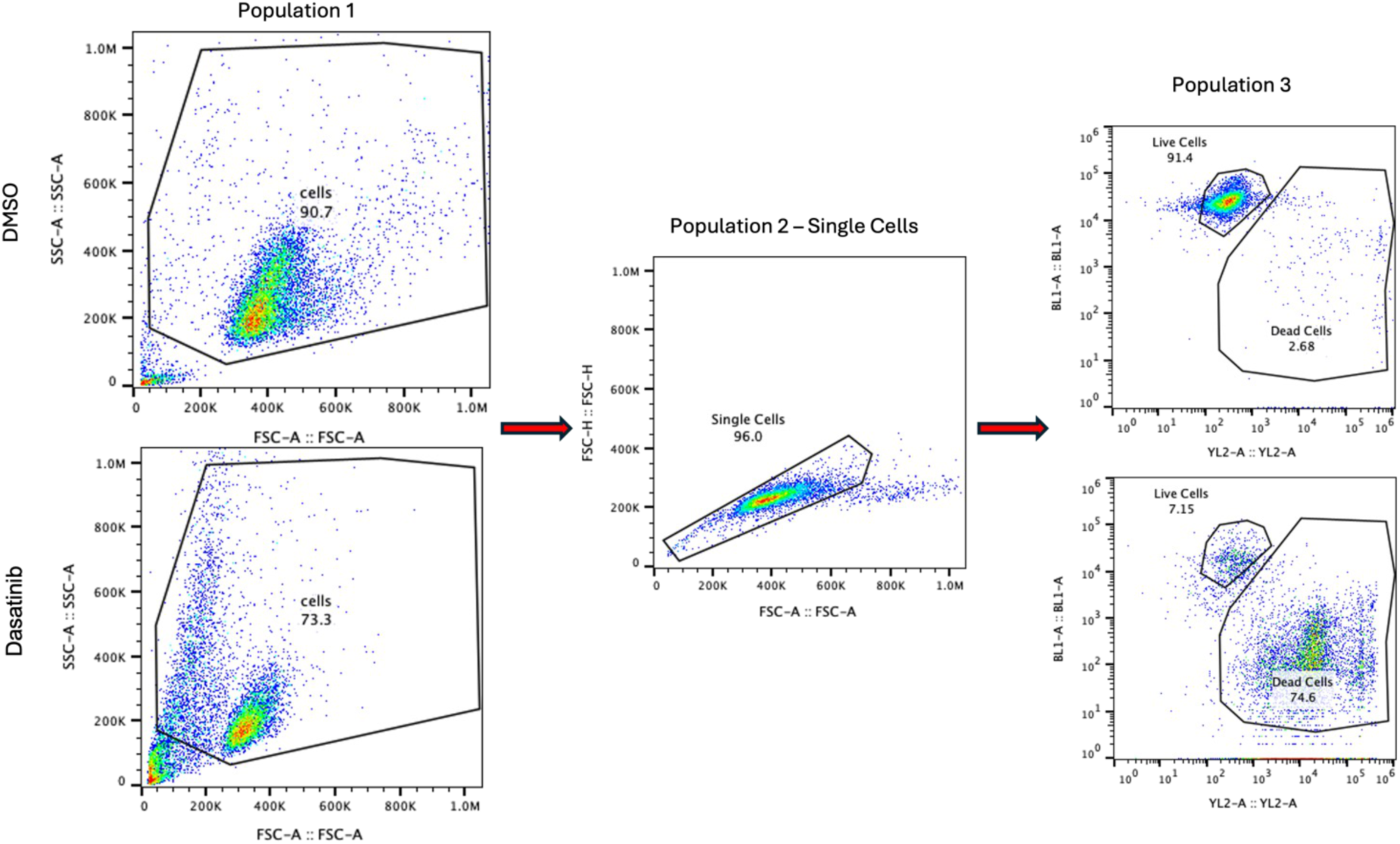
Flow cytometry gating strategy Gating strategy for flow cytometry data of k562/CRISPRi cells to study the effect of dasatinib on CRISPRi constructs targeted against HIC-2, PTH2R, and ZFPM2 stained with 1ug/ml EthD-1 and 2 μM calcein-AM analyzed by flow cytometry. The first panel shows all events with a gate (population 1) for cells. The second panel shows population 1 with a gate for singlets (population 2). The third panel shows Quarter 1 = Green or BL1-A for live cells and Quarter 3 = Red or YL2-A for dead cells (population 3).

## Notes

### Summary of Updates

gRNA cloning method listed an outdated version of the plasmid backbone. This information was updated. The revised plasmid backbone will be available upon request and will be available from addgene upon publication of the manuscript.

