## Supplemental Text 1 for "Simultaneous single-cell CRISPR, RNA, and ATAC-seq enables multiomic CRISPR screens to identify gene regulatory relationships"

**CAT-ATAC Protocol**

Before starting:

- Set heat block to 95°C for incubation of digitonin
- Cool swing-bucket centrifuge to 4°C

1. Anneal oligos 1 and 2 to make the RT splint duplex according to Oligo Annealing protocol by IDT. Store in -20°C.
2. Extract nuclei according to 10x protocol for cell lines.
3. Transpose nuclei according to 10x multiome protocol.
4. At **GEM Generation & Barcoding step (2.1)**, add spike-in RT primer (diluted to 0.5uM) and Hi-T4 ligase (NEB, M2622S).

| | For 1x (ul) |
|---|---|
| Barcoding reagent mix | 49.5 |
| Template switch oligo | 1.1 |
| Reducing agent | 1.9 |
| RT-splint duplex, 0.5uM | 1 |
| Barcoding enzyme mix | 7.5 |
| Hi-T4 DNA ligase | 3 |
| Total | 64 |

5. Incubate GEMs with the following conditions:

| GEM Incubation | | |
|---|---|---|
| Step | Temperature | Time |
| 1 | 25°C | 00:30:00 |
| 2 | 53°C | 00:45:00 |
| 3 | 25°C | 00:30:00 |
| 6 | 4°C | Hold |

6. Follow 10x protocol until **Post GEM Incubation Cleanup – SPRIselect step (3.2)**. Instead of 1.8x SPRI beads cleanup, use 3.2x SPRI beads (160ul) and 1.8x isopropanol (90ul) to ensure single-stranded cDNA from sgRNAs can be purified.
7. Make ATAC and GEX libraries as instructed in 10x protocol.
8. For CRISPR capture library:
   i. Take 40ul pre-amp product, perform 0.6x-1.0x bead selection to remove large cDNA fraction. Add 60ul EB to make total volume 100ul. First beads addition is 60ul, second beads addition 40ul. Elute in 30ul.
   ii. Perform Feature PCR using primers 8 and 5, 5ul template, in a 100ul reaction volume with Kapa HiFi HotStart ReadyMix (Roche, KK2601).

| Feature PCR | | |
|---|---|---|
| | For 1x (ul) | x2.2 |
| HiFi Master Mix | 50 | 110 |
| 10uM Forward primer (oligo 8) | 4 | 8.8 |
| 10uM Reverse primer (oligo 5) | 4 | 8.8 |
| | 58 | |
| DNA | 5 | |
| Water | 37 | 81.4 |
| Total | 100 | |

| Feature PCR Cycling (10 cycles) | | |
|---|---|---|
| Step | Temperature | Time |
| 1 | 98$^{o}$C | 00:00:45 |
| 2 | 98$^{o}$C | 00:00:20 |
| 3 | 58$^{o}$C | 00:00:05 |
| 4 | 72$^{o}$C | 00:00:05<br>Go to step 2, repeat 9x |
| 5 | 72$^{o}$C | 00:01:00 |
| 6 | 4$^{o}$C | Hold |

i. Use 2ul to run on Tapestation D1000 HS. Check if amplicon size is within expected range.

ii. Beads cleanup with 0.8x-0.9x SPRI. Elute in 30ul. Confirm size of purified product with Tapestation D1000 HS again. Example TS trace is shown below.

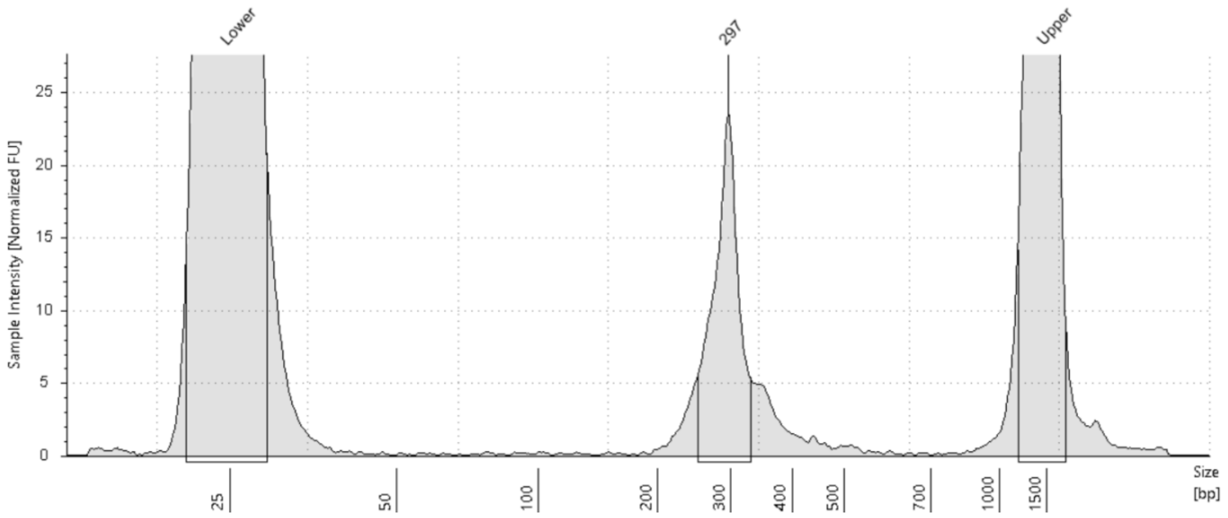


iii. Perform Sample Index PCR with primers 8 and 6a-f, 5ul template, in a 100ul reaction volume.

| Sample Index PCR |
|---|

| | For 1x (ul) | x2.2 |
|---|---|---|
| HiFi Master Mix | 50 | 110 |
| 10uM Forward primer (oligo 8) | 4 | 8.8 |
| 10uM Reverse primer (oligo 6a-f) | 4 | 8.8 |
| | 58 | |
| DNA | 5 | |
| Water | 37 | 81.4 |
| Total | 100 | |

| Sample Index PCR Cycling (9 cycles) | | |
|---|---|---|
| Step | Temperature | Time |
| 1 | 98$^{o}$C | 00:00:45 |
| 2 | 98$^{o}$C | 00:00:20 |
| 3 | 60$^{o}$C | 00:00:30 |
| 4 | 72$^{o}$C | 00:00:20<br>Go to step 2, repeat 8x |
| 5 | 72$^{o}$C | 00:01:00 |
| 6 | 4$^{o}$C | Hold |

i. Use 1ul to run on Tapestation D1000.
ii. Clean up with 0.7x-0.8x beads for double-sided size selection. Elute in 30ul. Use 1ul to run on Tapestation D1000 to confirm size. Example TS trace is shown below.

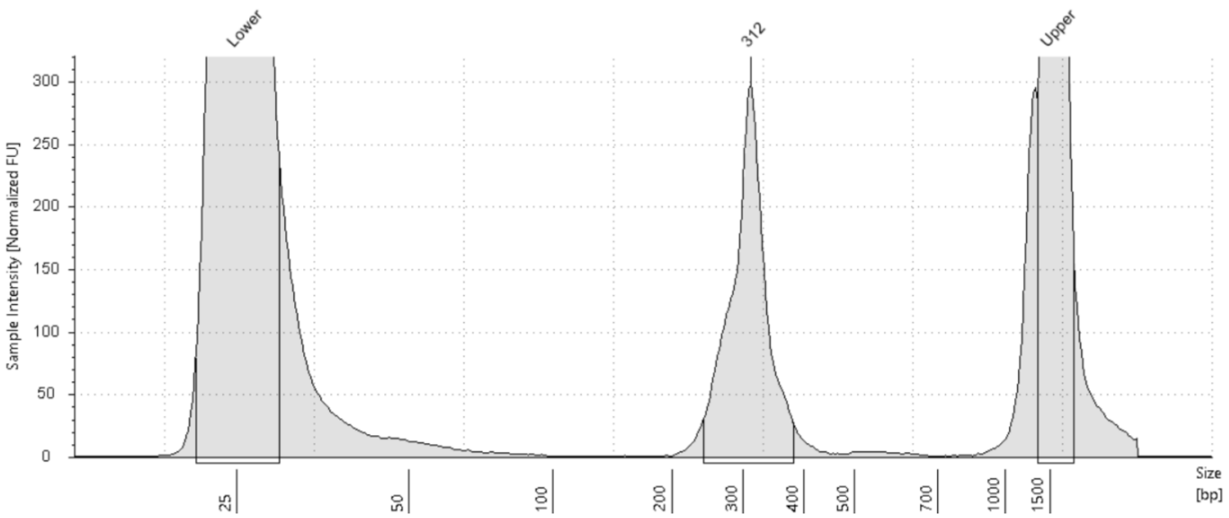


**Supplementary Notes:**

1. Structure of the CRISPR library is shown below:

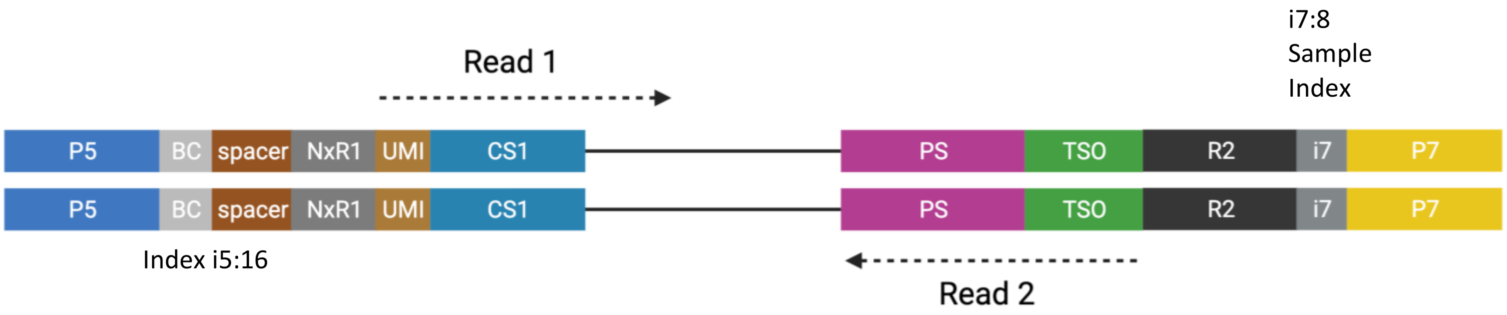


2. CRISPR library should be pooled with ATAC library and sequenced together. Sequencing configuration should be 151x8x24x151 (at least 75bp PE to be able to sequence through protospacer region). Aim for 25,000 read pairs per nucleus for ATAC library, and 5,000 read pairs per nucleus for CRISPR library.

3. Oligonucleotide sequences

| Name | Description | Sequence |
|---|---|---|
| 1 | RT primer CS1_12bp UMI | /5Phos/TCGTCGGCAGCGTCAGATGTGTATAAGAGACAGNNNNNNNNNNNNTTGCTAGGACCGGCCTTAAAGC |
| 2 | splint with LNA | TGACGCTGCC+G+A+C+G+ACAGACGCG/3Phos/ |
| 5 | R2-TSO primer | GTGACTGGAGTTCAGACGTGTGCTCTTCCGATCTAAGCAGTGGTATCAACGCAGAG |
| 8 | Partial P5 forward | AATGATACGGCGACCACCGAGA |
| 6a | P7-i7-R2 primer A1 | caagcagaagacggcatacgagatTTCGCAGTgtgactggagttcagacgtgtgctcttccgatct |
| 6b | P7-i7-R2 primer A2 | caagcagaagacggcatacgagatCGAGACTAgtgactggagttcagacgtgtgctcttccgatct |
| 6c | P7-i7-R2 primer A3 | caagcagaagacggcatacgagatACAGCTCAgtgactggagttcagacgtgtgctcttccgatct |
| 6d | P7-i7-R2 primer B1 | caagcagaagacggcatacgagatAAGTGTCGgtgactggagttcagacgtgtgctcttccgatct |
| 6e | P7-i7-R2 primer A1_1 | caagcagaagacggcatacgagatAGTAAACCgtgactggagttcagacgtgtgctcttccgatct |
| 6f | P7-i7-R2 primer A1_2 | caagcagaagacggcatacgagatCCGTTTAGgtgactggagttcagacgtgtgctcttccgatct |

4. CRISPR library index sequences

| Primer name | i7 index | i7 for sample sheet |
|---|---|---|
| 6a | TTCGCAGT | ACTGCGAA |
| 6b | CGAGACTA | TAGTCTCG |
| 6c | ACAGCTCA | TGAGCTGT |
| 6d | AAGTGTCG | CGACACTT |
| 6e | AGTAAACC | GGTTTACT |
| 6f | CCGTTTAG | CTAAACGG |

5. Note: Do not use the following ATAC indexes from the multiome kit to avoid index clashing with CRISPR library indexes (hamming distance <2):

SI-NA-B1
SI-NA-A5
SI-NA-B6
SI-NA-G11
SI-NA-C11
